# Construction and characterization of a genome-scale ordered mutant collection of *Bacteroides thetaiotaomicron*

**DOI:** 10.1101/2021.12.27.474270

**Authors:** Heidi A. Arjes, Jiawei Sun, Hualan Liu, Taylor H. Nguyen, Rebecca N. Culver, Arianna I. Celis, Sophie Jean Walton, Kimberly S. Vasquez, Feiqiao Brian Yu, Katherine S. Xue, Daniel Newton, Ricardo Zermeno, Meredith Weglarz, Adam Deutschbauer, Kerwyn Casey Huang, Anthony L. Shiver

## Abstract

Genomic analyses have revealed how the gut microbiota impacts human health. However, knowledge about the physiology of most gut commensals is largely lacking. Here, we sorted cells from a pooled library to construct an ordered collection of transposon-insertion mutants in the model commensal *Bacteroides thetaiotaomicron*. We applied a pooling strategy with barcode sequencing to locate mutants and created a condensed collection with single insertions in 2,565 genes. This effort enabled the development of an accurate model for progenitor-collection assembly, which identified strain-abundance biases and multi-insertion strains as important factors that limit coverage. To demonstrate the potential for phenotypic screening, we analyzed growth dynamics and morphology of the condensed collection and identified growth defects and altered cell shape in the sphingolipid-synthesis gene BT0870 and the thiamine scavenging gene BT2397. Analyses of this collection and utilization of the platform described herein to construct future ordered libraries will increase understanding of gut commensal physiology and colonization strategies.

## Introduction

The gut microbiota is a complex community that plays a pivotal role in digestion, colonization resistance, immune signaling, and other health outcomes (1, 2). The mechanisms by which the members of our microbiota exert their effect remain largely unknown. Tools and resources for genetic analysis of microbiota members could be transformative for the mechanistic investigation of microbe-host interactions, but they must be generalizable and scalable to the diverse members of this critical microbial community. The Bacteroidetes phylum includes many species that are prevalent in mammalian gut microbiotas (3) and play important roles in human health (4-7). *Bacteroides thetaiotaomicron* (commonly referred to as *B. theta*) is a model organism for the *Bacteroides* genus and is of particular interest due to its ability to metabolize complex polysaccharides (8-11) and its capsule production (12, 13), enabling survival within many environmental niches in the gut (14, 15). *B. theta* antigens have been linked to host T-cell responses, which differ depending on diet (16), and *B. theta* exhibits several connections with host metabolism through lipid (17) and outer membrane production (18). However, genetic mechanisms underpinning the interactions of *B. theta* with its host remain challenging to study because of the effort and time required to generate targeted gene disruptions (19-21). The *B. theta* genome contains 4902 currently annotated genes.

Genetic disruption via transposon insertion has enabled the creation of pooled mutant libraries that have been used in combination with deep sequencing for genome-wide fitness assays in many species. In *B. theta*, a library of ∼35,000 transposon mutants facilitated genome-scale interrogation of genes important for *in vitro* growth and for *in vivo* colonization of the mouse gut using insertion sequencing (INSeq) (22, 23). The inclusion of a unique DNA barcode within each transposon transforms the relatively laborious and costly transposon insertion sequencing (TnSeq) protocol into a simple barcode amplification and sequencing protocol (BarSeq) once the mapping between barcode and insertion location is established (24). The resulting reduction in cost and effort dramatically enhances the throughput of pooled screens that map genotype to phenotype (25). We recently constructed a library of >300,000 barcoded transposon mutants in *B. theta* and characterized the fitness of this pooled library across hundreds of *in vitro* conditions and during colonization of germ-free mice (26). These pooled screens provided myriad functional and physiological insights, including specific phenotypes for 516 genes, and validation of mutant phenotypes was greatly facilitated by isolation of mutants of interest. Moreover, isolation of single strains is necessary to uncover mutants with phenotypes that are masked in a pooled population, for example involving secretion of a molecule that is shared among the entire population regardless of genotype (27-29). To facilitate isolation from a pooled library, we recently developed a protocol based on cell sorting that is effective for even strict anaerobes (30).

Here, we report the creation of a genome-scale, ordered collection of *B. theta* transposon mutants in a streamlined, efficient, and generalizable manner. We first arrayed a progenitor collection of nearly 30,000 potential mutants. We then applied a pooled sequencing strategy to identify the mutant in each well, and re-arrayed selected strains to create a condensed collection of 2,565 strains with a single transposon insertion per gene that covers 52% of the genome. Next, we used the statistics of our progenitor collection along with a quantitative model of the assembly process to identify the factors that reduce coverage during sorting, providing a template for future optimization. Finally, we characterized the growth dynamics and morphology of each mutant in the condensed library. We found that most mutants exhibited nearly wild-type growth and cell shape. Among the few outliers, we identified mutants in a gene involved in sphingolipid synthesis and a gene involved in thiamine scavenging that were defective in growth and for which a subpopulation of cells was elongated relative to wild type. Given the substantial time and effort required for the construction of targeted mutants in *B. theta*, our collection and strategy for assembly should serve as an impactful resource to gut microbiome research, particularly for investigating mechanisms of commensal survival and function in the mammalian gut.

## Results

### Construction of a 2,565-gene ordered transposon-insertion collection in *B. theta*

To assemble a community resource for high-throughput phenotypic screens in *B. theta,* we used a previously developed protocol based on fluorescence-activated cell sorting (FACS) (30) to isolate individual mutants of *B. theta* VPI-5482 from a pool of >300,000 randomly barcoded transposon insertions covering 4532 of the 4902 currently annotated genes of *B. theta* (26) (Figure 1). In a pilot experiment to evaluate the isolation efficacy of cell sorting, we isolated 3,493 strains in 40 96-well plates that included 2,075 strains with single insertions covering 1,029 unique genes (30). While this resource proved useful for certain follow-up mechanistic studies (26, 29), the absence of a large majority of non-essential genes motivated a more extensive ordered collection. In the present study, we first expanded the coverage of this progenitor collection by sorting mutants into an additional 262 96-well plates to create a 302-plate progenitor collection (Figure 1). We then used BarSeq with a plate-well pooling strategy to locate barcodes within the new progenitor collection (Figure 1, Methods).

**Figure 1:**
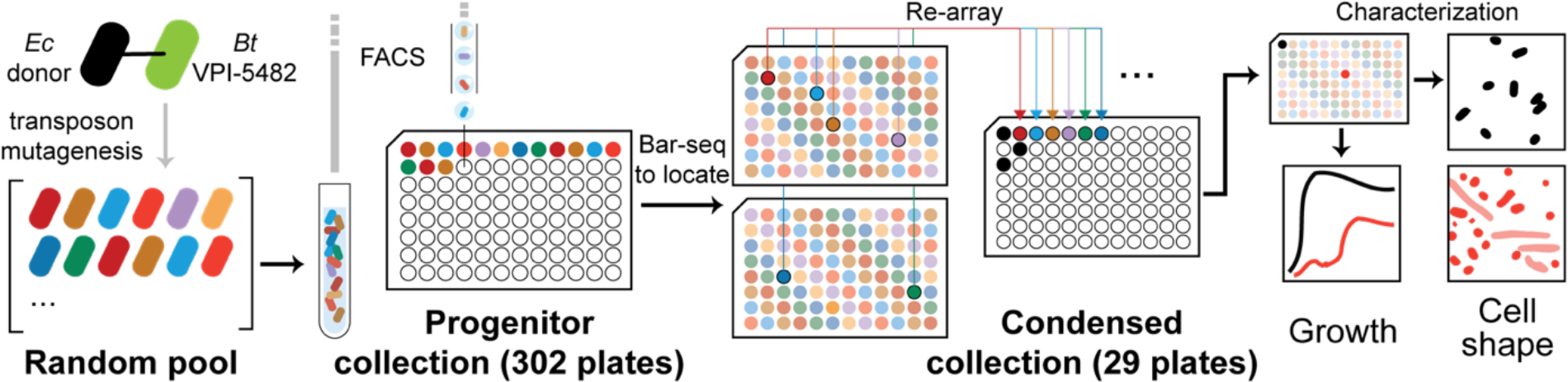
Workflow for the construction of an ordered collection of *B. thetaiotaomicron* VPI-5482 transposon mutants. Single cells from a random pool of >300,000 mutants (30) were sorted into 302 96-well plates using FACS to create a progenitor collection. After using barcode sequencing to locate strains within the progenitor collection, representative mutants for individual genes were re-arrayed to create a condensed collection of 29 96-well plates covering 2,565 genes. To demonstrate the screening potential of this collection, growth and cell shape phenotypes of each strain were characterized during growth in rich medium (BHIS).

In the 302-plate progenitor collection, 83% of barcodes could be located unambiguously in a single well of the collection, 4% could be located with a probability >0.85, and 13% of barcodes could not be located in the collection with high confidence (Methods). A total of 2,677 genes were covered by at least one single-insertion strain in the larger progenitor collection, more than doubling the coverage of the non-essential genome relative to our initial 40-plate isolation (Figure 2). This coverage value takes into account all single-insertion barcodes in the progenitor collection, regardless of potential contamination or location ambiguity. The progenitor collection covers 326 of 415 genes currently annotated as encoding carbohydrate active enzymes (http://www.cazy.org) (31) (Figure 2B), highlighting its potential for investigating the connection between carbohydrate utilization and host colonization along with many other phenomena.

**Figure 2:**
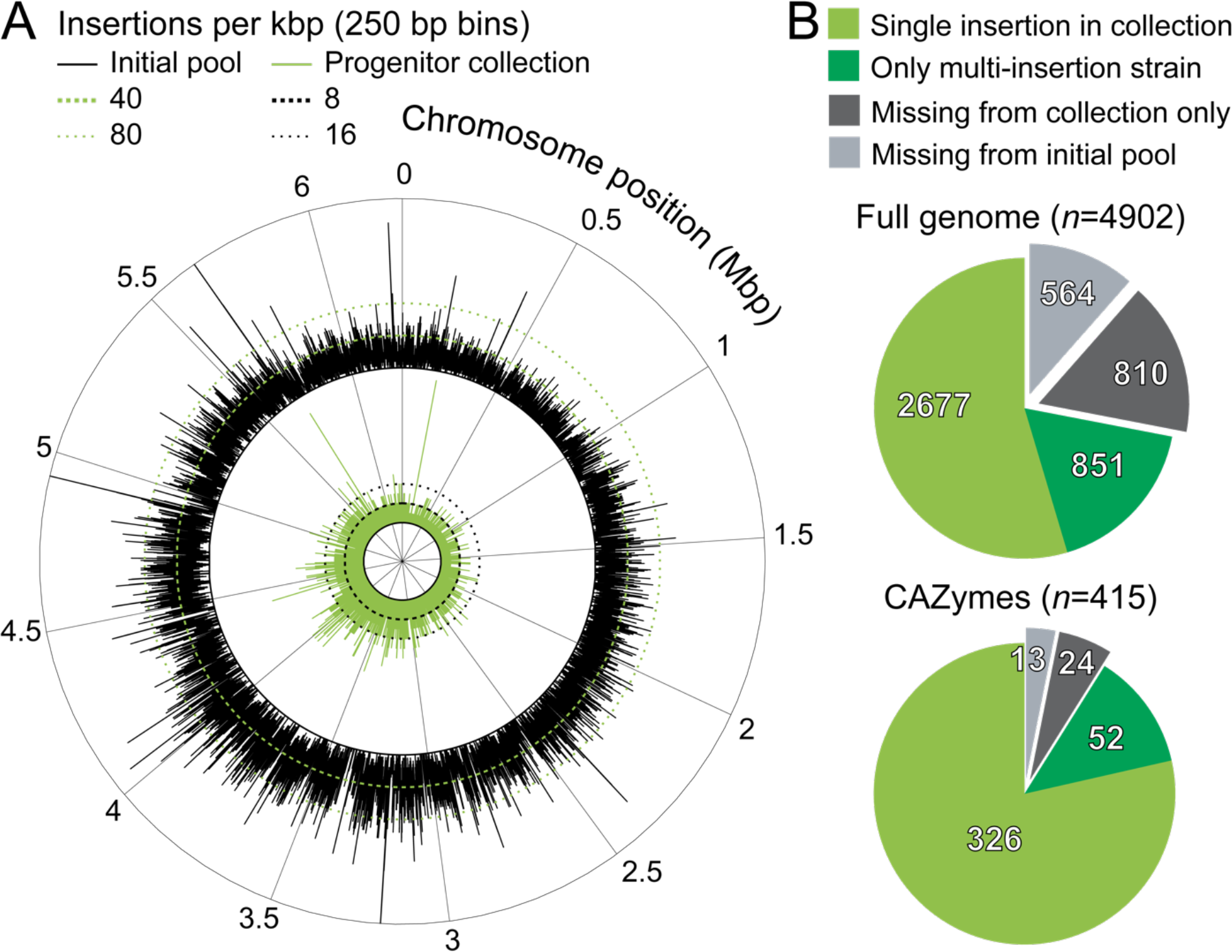
The progenitor collection covers a large fraction of the genome. A) Patterns of transposon insertion density across the chromosome were largely similar between the initial pool and the progenitor collection. The insertion density across the chromosome is plotted for the initial pool (∼150,000 insertions that map to the genome; outer circle, black) and the progenitor collection (∼12,000 insertions; inner circle, green). The number of barcodes detected in the initial pool used for sorting was smaller than the total detected in the original random pool, presumably due to loss of strains during passaging. Insertion density was calculated from the number of uniquely barcoded transposons in 250-bp bins and does not reflect the relative abundance of individual barcodes. Only single-insertion strains were considered when calculating insertion density for the progenitor collection. B) Coverage in the progenitor collection of the *B. theta* genome (top) and the carbohydrate active enzyme (CAZyme, bottom) functional category. Genes with single-insertion strains were candidates for inclusion in the condensed collection (light green). Some genes were only covered by strains with multiple insertions (dark green), and some had no representative insertions in the initial pool and thus were unlikely to be captured (light gray). Other genes were covered in the initial pool but not captured in the progenitor collection (dark gray); these genes are likely to be accessible with larger collection sizes or with alternative sorting parameters. *n* specifies the number of genes.

Our progenitor collection substantially expands the scope of gene functions covered relative to existing mutant collections of *B. theta*: our collection contains 1,866 unique genes, a previously described non-barcoded mutant collection (23) contains 811 unique genes, and 699 genes overlap between the two collections (Figure S1). However, the sheer size of the progenitor collection is prohibitive for screening and distribution. Thus, we re-arrayed isolates from the progenitor collection to assemble a condensed collection in which transposon disruptions of each gene were represented by one strain (Figure 1, Dataset S1). To simplify the interpretation of mutant phenotypes, we required that the barcoded insertion map between the first 5% and the last 10% of the open reading frame and only map to one location in the genome. Potentially contaminated wells were inferred from overlapping barcode locations that were predicted with high confidence (>0.85) and were excluded from re-arraying. When more than one barcoded insertion strain for the same gene was isolated in the progenitor collection, we prioritized the insertion closest to the middle of the gene. Our re-arraying reduced the size of the collection by an order of magnitude (from 302 to 29 96-well plates), facilitating high-throughput screening (Figure 1). Beyond the 2,565 genes covered by the condensed collection (Dataset S1), >100 additional genes could be accessible in the progenitor collection with future efforts to clarify potential contamination and disambiguate repeated barcodes. Furthermore, the progenitor collection covers 851 additional genes that are only represented by strains with multiple insertions (Dataset S2); these genes were not included in the condensed collection but are available as a resource. Each of the 29 96-well plates of the condensed collection included three empty wells, either to serve as a blank or to enable the inclusion of wild-type cells as a control in future screens.

### Progenitor collection reveals how bias in library composition and the presence of multi-insertion strains limits coverage

In an ideal progenitor collection, every available well would be occupied by a single strain with a potentially useful mutation. In practice, there are inefficiencies in assembling the collection that reduce the fraction of wells for which a useful barcode is recovered. Thus, this fraction (which we refer to as the assembly efficiency) is an important factor to consider when designing a progenitor collection to improve the odds that the collection approaches saturation of the initial pool. Given the coverage of the condensed collection (52% of annotated genes), we were interested in determining the progenitor-collection size that would have allowed us to approach saturation of the initial pool (which contains 88.3% of annotated genes), the impact of assembly efficiency, and the factors that most influenced assembly efficiency. The large size of our progenitor collection allowed us to address these outstanding questions, thereby informing and improving future efforts. We modeled the process of assembling the *B. theta* progenitor collection to identify the factors that most affected its genome coverage. To simplify our model, we focused on the 262 96-well plates sorted in this study, as all transposon mutants originated from the same pooled library culture.

Leveraging the accurate quantification of strain abundance in the initial pool available through BarSeq, we used a Monte Carlo approach (repeated random sampling) (32) to simulate the isolation of barcodes from the initial pool and calculated the genome coverage (number of genes represented by ≥1 insertions) across a range of progenitor collection sizes with *b* isolated barcoded strains (Figure 3A). To accurately model the saturation curve, we needed to account for inefficiencies in library creation and sorting that reduced the number of useful barcoded strains isolated in the progenitor collection (Figure 3A). We identified three such factors and estimated their effect size from the final statistics of the 262-plate collection. First, 13% of wells did not contain a barcode and 2% contained >1 barcode. Accounting for these two factors, the fraction of wells with a single barcode (*b*/*w*) is 0.85. Second, 13% of barcodes did not map to the genome (*f*mapped=0.87) and, of the mapped barcodes, 32% were associated with more than one insertion site (*f*single=0.68). The product of *b*/*w*, *f*mapped, and *f*single is an estimate of the overall assembly efficiency *K*=0.504 (Figure 3A).

**Figure 3:**
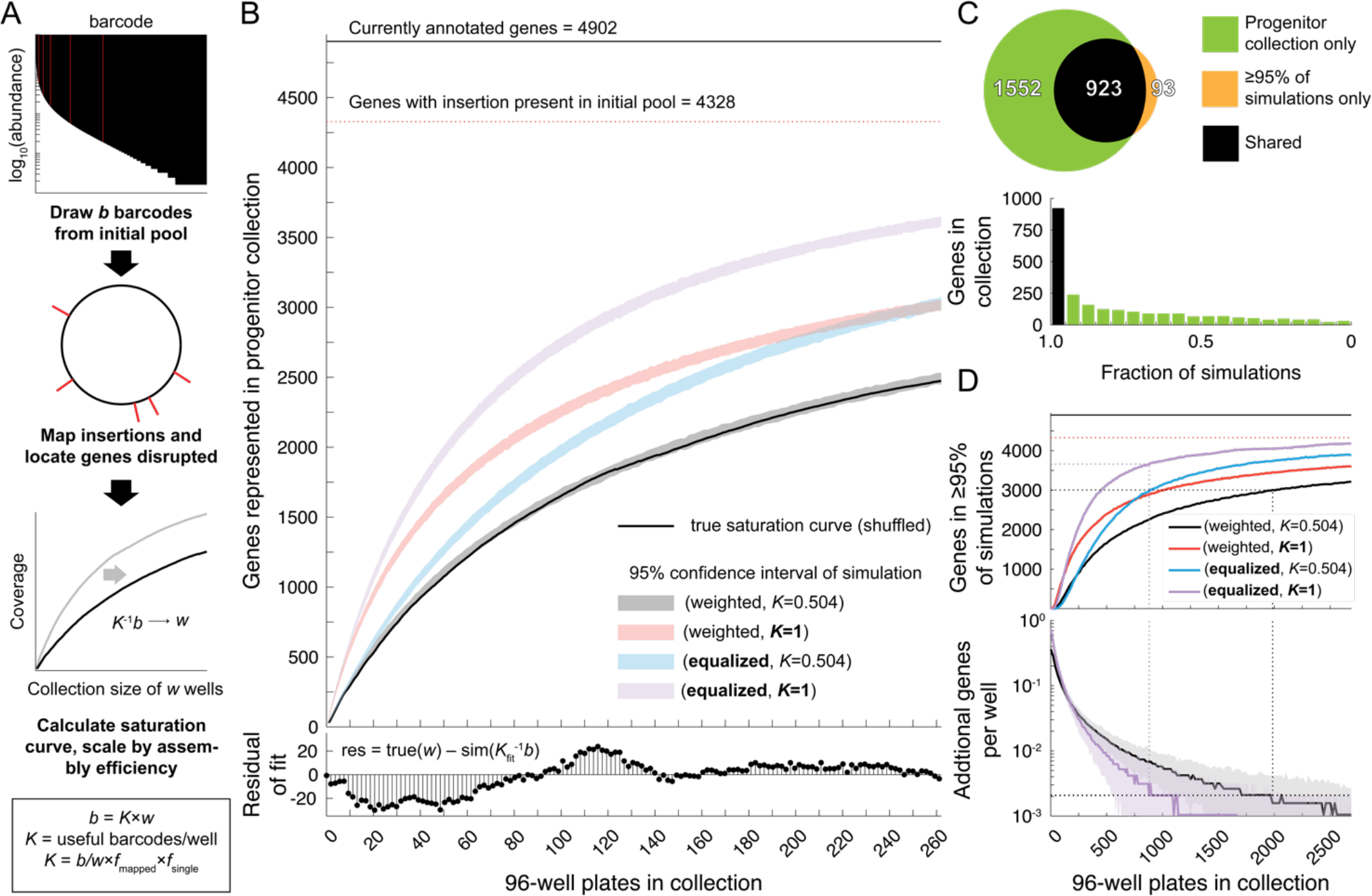
Monte Carlo simulations reveal factors that limit genome coverage in the progenitor collection. A) Workflow for simulating genome coverage in the progenitor collection. BarSeq was used to estimate the identity and abundance of insertions in the initial pool used for sorting. Relative barcode abundance was used as a probabilistic weight to draw *b* barcodes from the initial pool. The insertions associated with each barcode were located on the genome and compared to the genome annotation to determine the coverage (the number of genes disrupted) of the simulation. To model the coverage of a collection of *w* wells, we estimated the assembly efficiency (*K*, the fraction of wells with a useful barcode): we counted the average number of barcodes per well, then factored in the requirements that the barcode had to be associated with an insertion at a defined genome location and had to be a single insertion. We scaled the collection size in the simulations by *K*^-1^ to account for assembly efficiency. B) (Top) The experimental saturation curve (black) is plotted along with 95% confidence intervals of simulated saturation curves. The total number of genes (4,902) and the total number of genes represented covered by the initial pool (4,328) are shown as black and dashed red lines, respectively. A saturated collection would have coverage that approaches the number of genes in the initial pool. Using the initial pool composition determined by BarSeq and accounting for the assembly efficiency of the collection allowed us to accurately model the true saturation curve (weighted, *K*=0.504). Hypothetical collections with ideal assembly efficiency (*K*=1), unbiased initial pools (equalized), or both resulted in collections with higher coverage. (Bottom) Fit residuals between the true collection (true(*w*)) and the simulated saturation curve sim(*K*fit *b*) obtained by fitting *K* to the true saturation curve (*K*_fit_=0.495). The simulation largely matched the experimentally measured saturation curve. C) (Top) The overlap between the genes covered in the 262-plate collection and the genes predicted to occur in ≥95% of 1,000 simulations of the same size is high. While the simulated gene set was largely represented in the 262-plate collection, the 262-plate collection contains many unique genes. (Bottom) Many genes in the collection were present in <95% of simulations (green bars), indicating that the 262-plate collection has not reached saturation of the initial pool. D) Simulations of larger collection sizes predict the requirements for reaching saturation. (Top) The number of genes that occur in ≥95% of 250 simulations is predicted to saturate within 2,679 plates only with both ideal assembly efficiency and equalized barcode abundance (equalized, *K*=1; purple line). The horizontal red dashed line represents the total number of genes covered by the initial pool, the saturation limit of any ordered collection. The black and purple dashed lines track collection size statistics for the original (weighted, *K*=0.504) and ideal (equalized, *K*=1) simulations, respectively, with a heuristic practical limit on collection size, as described below. (Bottom) The incremental increase in coverage per well added to the collection can be used to guide the practical size limit of progenitor collections. For example, at a hypothetical incremental efficiency of <2×10^-3^ (horizontal dashed line), an additional 10 96-well plates will only isolate 1–2 additional genes. The collection sizes of both the original (weighted, *K*=0.504; black dashed line) and ideal (equalized, *K*=1; purple dashed line) simulation are represented as vertical dashed lines.

We accurately predicted the experimentally measured saturation curve of the 262-plate progenitor collection by scaling the collection size of the simulated saturation curves by *K*^-1^ (Figure 3B, top). Fitting simulation results to the measured saturation curve resulted in a similar estimate of *K* (0.495) with low bias in the residuals of fit throughout the saturation curve (Figure 3B, bottom), suggesting that scaling of the simulated saturation curve can accurately model the assembly of a progenitor collection.

We next asked whether we could predict the composition of the progenitor collection in addition to the saturation curve. We performed multiple simulations and quantified the fraction in which a representative insertion strain for each gene was included. While 91% of genes predicted to be present at high confidence (≥95% of simulations) were indeed isolated in the 262-plate collection, 63% of the progenitor collection was composed of genes whose probability of being included in the simulation was estimated to be low (<95% of simulations) (Figure 3C). These data suggest that the 262-plate collection has not reached saturation, and motivate a systematic assessment of the requirements for achieving saturation, for this and future progenitor collections.

### Prediction of design requirements for achieving saturation of coverage

Collectively, the results above highlight two outstanding factors limiting coverage. First, the largest individual factor influencing assembly efficiency *K* was the number of barcodes that mapped to multiple insertion sites (1-*f*single). Second, a considerable fraction of wells (19%) in the 262-plate progenitor collection were wasted on repeats of previous barcode isolations, presumably due to biases in strain abundances in the initial pool. To quantify the potential increase in coverage from overcoming these technical challenges, we simulated the assembly of progenitor collections with *K*=1 and/or equalized strain abundance. The 262-plate collection would have covered 539 or 566 more genes from the initial pool after resolving low assembly efficiency or strain abundance biases, respectively (Figure 3B, top). Resolving both issues would have resulted in the 262-plate collection covering 1,138 more genes from the initial pool (Figure 3B, top).

Finally, we estimated the effort that would be required to assemble a truly saturated collection and the effect of assembly efficiency and bias in strain abundance on this outcome. We simulated the creation of progenitor collections of up to 2,679 96-well plates (the number that would completely fill a standard -80 °C freezer) and quantified the number of genes present in ≥95% of simulations (Figure 3D). After sorting 2,679 plates, the initial pool would have covered 3,213 of the 4,328 genes in the pooled library with high confidence (Figure 3D). Equalizing strain abundance, optimizing assembly efficiency (*f*single=*f*mapped=*b*/*w*=1 so that *K*=1), or both reduced the number of 96-well plates required to reach this coverage to 1,092, 1,351, and 551, respectively (Figure 3D). These data indicate that the initial pool limits the practicality of achieving a truly saturated library and that both technical challenges will need to be overcome before libraries that fully saturate the initial pool can be assembled with reasonable effort.

For a more practical estimate of the coverage that could be achieved, we tracked the incremental efficiency of isolating additional gene disruptions after adding wells to a progenitor collection of a given size. When the incremental efficiency drops below 2×10^- 3^, adding 10 96-well plates to the progenitor collection is likely to isolate only 1–2 additional genes. Using this benchmark to constrain the practical size of a progenitor collection, we found that our initial pool could be used to create a 1,985-plate collection covering 3,002 genes with high confidence before isolating additional genes would become impractical (Figure 3D). By contrast, a library with equalized strain abundance and *K*=1 would cover 3,659 genes with high confidence in an 881-plate library before incremental efficiency decreased below 2×10^-3^ (Figure 3D). Future studies will benefit from initial pools with strain abundances that are as uniformly distributed as possible.

### Characterization of growth dynamics reveals a small subset of mutants with inhibited growth

Previous studies of genome-scale knockout libraries in *E. coli* revealed that only a small subset of gene deletions perturb growth dynamics in rich media (33, 34). Likewise, *E. coli* cell shape is unaltered by most gene deletions (33, 35, 36). Nonetheless, unbiased genome-wide screens have identified important components of the pathways controlling growth and cell shape (36, 37). Given the paucity of similar investigations in most other organisms, our condensed collection provided an exciting resource for discovery regarding *B. theta* physiology. To this end, we measured the growth dynamics of the condensed collection in a rich medium (BHIS) and used high-throughput imaging (38) to characterize cell morphology of each strain in stationary phase (Figure 4A).

**Figure 4:**
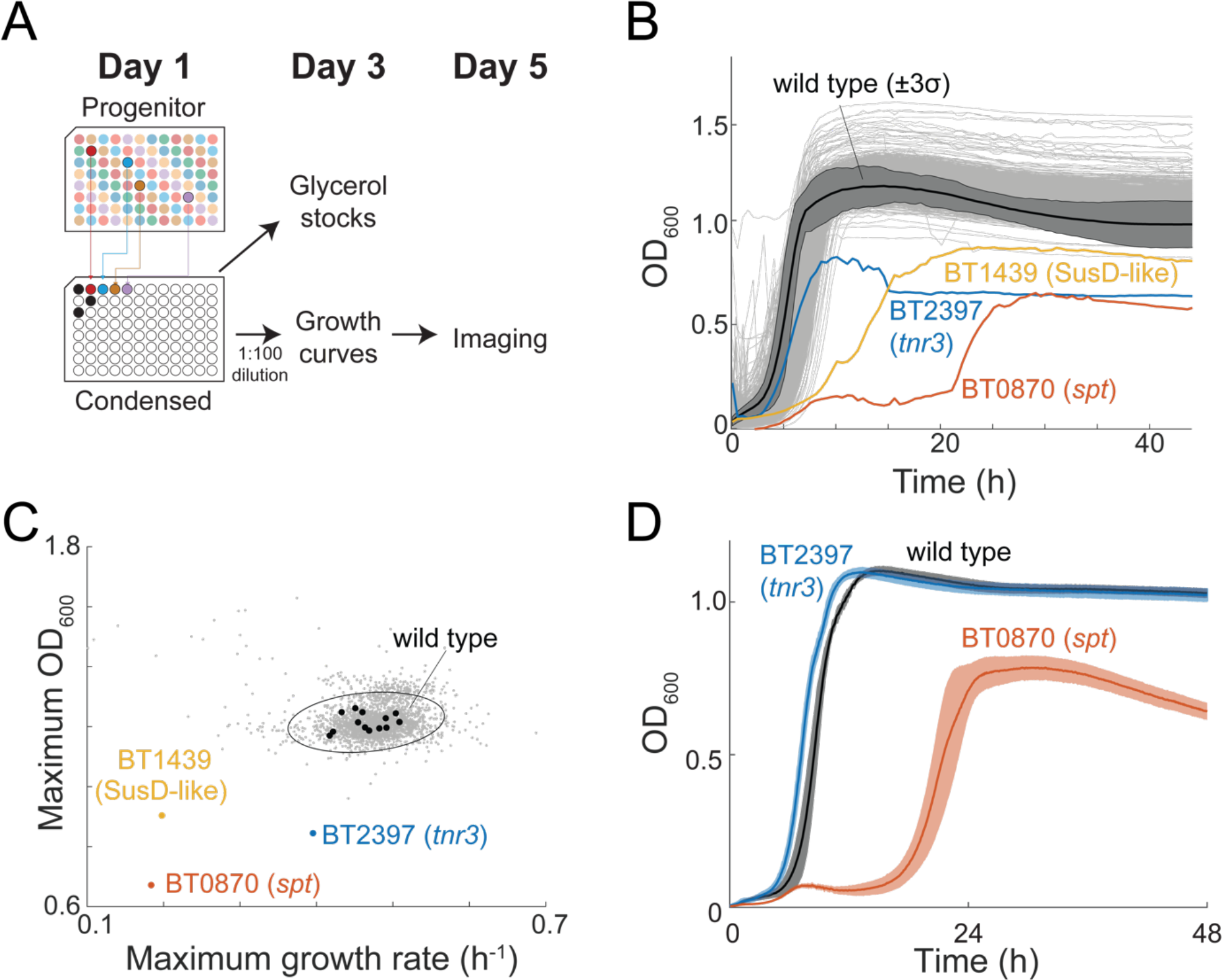
High-throughput screening reveals a small set of mutants with growth defects. A) Schematic of the pipeline for measuring growth and cell shape phenotypes of the mutants in the condensed collection. B) Growth in BHIS was similar to wild-type for the vast majority of mutants (gray curves). BT0870 (*spt*, sphingolipid synthesis, red), BT2397 (*tnr3*, thiamine scavenging, blue), and BT1439 (SusD-like, yellow) mutants exhibited growth defects. The average of all wild-type growth curves is shown in black with the dark gray shaded region representing 3 standard deviations. C) Most mutants exhibited similar maximum growth rate and maximum OD600 as wild type. Wild type replicates (black circles) exhibited a similar spread as most of the mutants. All mutants except the outliers described in (B) are shown in gray. The oval indicates 3 times the standard deviations and covariance. D) After passaging through a colony, the growth phenotype of the BT0870 (*spt*) mutant from the progenitor collection was maintained, while the BT2397 (*tnr3*) mutant reverted to approximately wild-type growth. Solid lines are averages and shaded regions indicate 1 standard deviation of *n*=6 biological replicates.

After re-arraying strains into the condensed collection, we diluted the 48-h cultures and regrew each strain in 96-well plates (Figure 4A). Wild-type cultures were added to empty wells in the condensed collection. Wild-type growth curves were reproducible across plates, with a maximum growth rate of 0.47±0.03 h^-1^ (1 standard deviation) that was reached ∼4 h after reinoculation and a maximum OD600 of 1.32±0.03 (1 standard deviation) reached after ∼13 h (Figure 4B,C). Most of the 2,565 mutant growth curves were similar to wild type (Figure 4B), with maximum growth rates and maximum OD600 within 3 standard deviations of the mean of the wild-type distribution (Figure 4C). In addition to the consistency of wild-type growth across plates, sets of technical replicates of each mutant growth curve identified the same outliers with low growth rates and yield, demonstrating that growth behaviors could be effectively compared between 96-well plates (Figure S2). Although a small fraction of mutants appeared to reach higher OD600 values than wild type, these high-yield outliers were not reproduced in the other replicate and may be due to technical imperfections in a minority of assayed wells (Figure S2D).

A small number of mutants had obvious growth phenotypes. The strongest growth defect was associated with an insertion in BT0870, which grew very slowly for ∼20 h before transitioning to rapid growth; these growth dynamics were preserved during liquid regrowth from a colony and during further liquid passaging (Figure S3, Figure 4D), suggesting that the later rapid growth is not due to a suppressor mutation. BT0870 (*spt*) encodes a serine palmitoyl transferase essential for the synthesis of sphingolipids in *B. theta* (39). An insertion in BT2397 had a similar maximum growth rate to wild type but reached a lower maximum OD600. BT2397 (*tnr3*) encodes a proposed thiamine pyrophosphokinase involved in scavenging thiamine from the environment (40). Finally, an insertion in BT1439 had a lower maximum growth rate and did not reach its maximum OD600 until 20 h. BT1439 encodes a SusD-like outer membrane protein important during mono-colonization of the murine gut and for the uptake of vancomycin (26). These results highlight the potential for discovery of fitness defects that would not be apparent in pooled screens.

### High-throughput imaging reveals morphological phenotypes in strains with growth defects

Using our high-throughput imaging protocol (38), we collected >140,000 images encompassing thousands of single cells for each strain in the condensed collection at the end of the 48-h growth curves in BHIS and used an automated computational pipeline (Methods) to segment each cell. During the transition to stationary phase, *B. theta* cells shorten and become rounder (Figure 5A). As with the growth curves, most strains in the collection had quantitatively similar stationary-phase cell morphology to wild type (Figure 5B). Nonetheless, the BT2397 (*tnr3*) mutant exhibited larger average cell width and cell length (Figure 5B), and the BT2397 (*tnr3*) and BT0870 (*spt*) mutants both exhibited increased heterogeneity in cell length and/or width. Thus, we focused on these two mutants for further characterization. In the BT0870 (*spt*) insertion, the mode of the distribution of cell areas was shifted to smaller values than wild type, and some cells were elongated (Figure 5C). These cell morphology defects are generally consistent with the known role of *spt* in membrane biosynthesis (39) and of fatty acid availability in bacterial cell-size determination (41).

**Figure 5:**
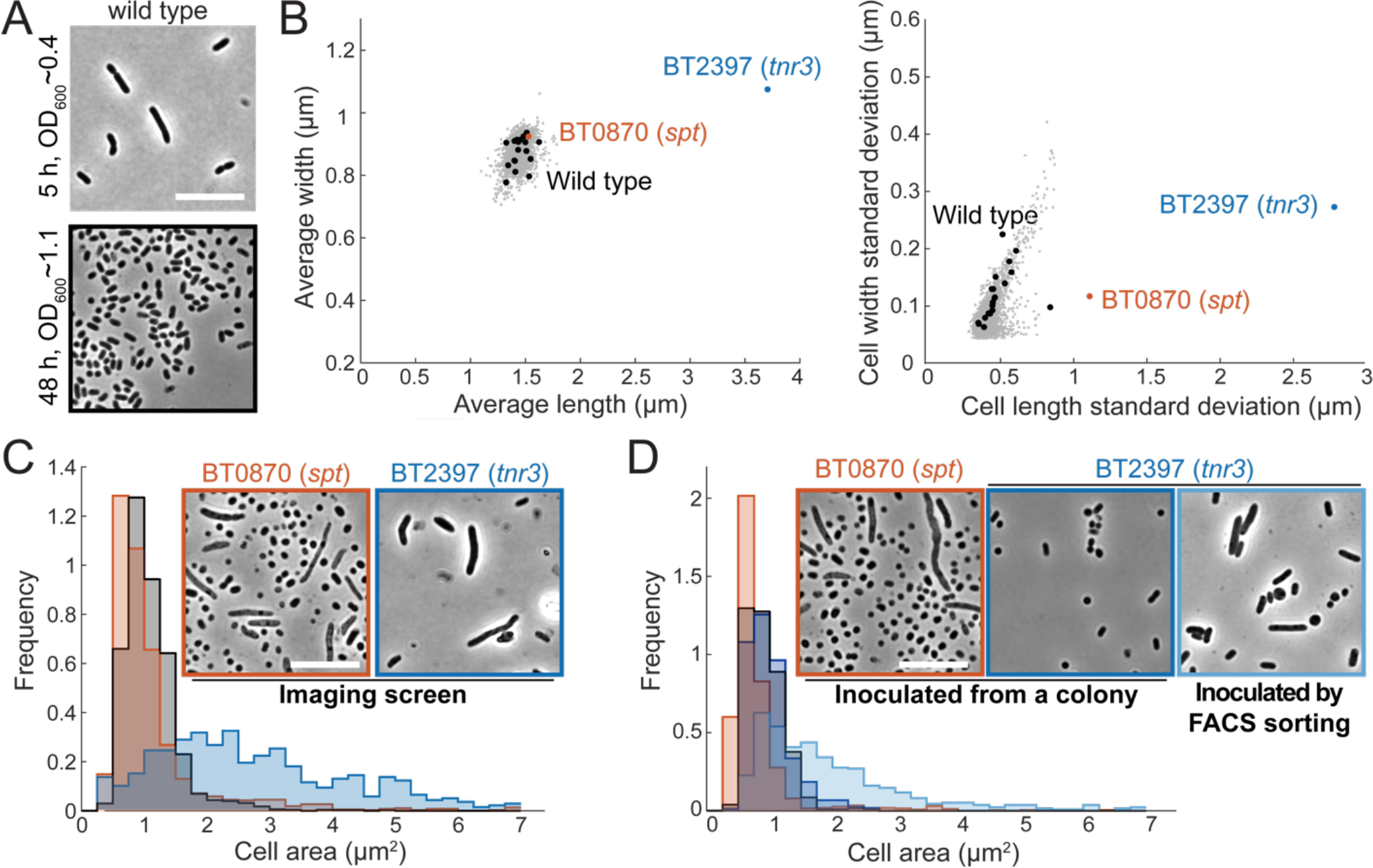
Morphological screening reveals filamentation phenotypes of the BT0870 (*spt*) and BT2397 (*tnr3*) mutants. A) Wild-type cells become shorter and rounder in stationary phase. Phase-contrast images of log-phase cells (OD600∼0.4) and stationary-phase cells (OD600∼1.1). Scale bar: 10 µm. B) (Left) BT2397 (*tnr3*) mutant cells had higher average length and width than wild-type cells. (Right) BT0870 (*spt*) mutant cells exhibited higher variation in cell length than wild type. All strains except these two outliers are shown in gray, and wild-type replicates are shown in black*. n*>1500 cells per strain. C) Morphological phenotypes were identified from screening. (Left) A subpopulation of BT2397 (*tnr3*) mutant cells were longer and subpopulations of BT0870 mutant cells were longer or shorter relative to wild-type cells. *n*>1500 cells per strain. (Right) Representative phase-contrast images of each mutant. Scale bar: 10 µm. D) Morphological phenotypes were retained after single cell isolation. BT2397 (*tnr3*) mutant cells were no longer elongated after passaging through a colony, but the filamentous phenotype was typically maintained after inoculating individual mutant cells into liquid BHIS using FACS with a gate to select for cells larger than wild type and culturing for 48 h. One representative culture of each strain/condition is shown and the inset shows a representative image of each culture. *n*>400 cells per strain.

### BT2397 (*tnr3*) mutant phenotypes require liquid culturing

The large cell sizes of the BT2397 (*tnr3*) insertion strain were surprising, given that mutants in genes related to the synthesis of thiamine pyrophosphate (BT0652) and the import of thiamine (BT2390, BT2396) were present in the condensed collection but did not display any growth or cell morphology defects. The morphological phenotype of this strain is unlikely to be due to an unidentified second-site mutation, as an independent mutant from the progenitor collection with an insertion in BT2397 (*tnr3*) shared the same elongated cell morphology (Figure S4A). The BT0870 (*spt*) mutant retained its growth and shape phenotype after passaging through a colony and regrowth in liquid (Figure 4D, 5D). The BT2397 (*tnr3*) mutant retained its shape phenotypes during liquid passaging from the progenitor stock (Figure 5C,D, S4), but its growth (Figure 4D) and stationary phase cell morphology (Figure 5D) phenotypes were lost upon liquid regrowth after passaging through a colony.

To investigate this difference in BT2397 (*tnr3*) outcomes further, we first evaluated the possibility that a mixed population of genetic backgrounds was present in the cryostock and that only the background with wild-type cell shape was stochastically recovered in the small number of colonies isolated. We isolated 64 single colonies after streaking a culture grown from the cryostock on a plate, recovered the colonies in BHIS broth, and measured growth curves and stationary-phase cell shape. All 64 cultures exhibited wild-type cell morphology (Figure S4B). To test whether growth as a colony was responsible for the reversion of the phenotypes, we used FACS to isolate 146 BT2397 (*tnr3*) single cells directly into BHIS broth and grew cultures from them. Eighty-three of these cells were isolated using a gate that selected for mutant cells with approximately wild-type-like size. The other 63 cells were isolated using a gate that selected for mutant cells larger than the average size of the BT2397 (*tnr3*) population. We re-passaged these 146 cultures in BHIS and measured stationary-phase cell morphology. Many of the large-gate and the small-gate cultures exhibited a substantial subpopulation of elongated cells (Figure S4B), matching our observations from the condensed collection (Figure 5D). These results indicate that the BT2397 (*tnr3*) mutant indeed has an elongated and heterogeneous shape phenotype in stationary phase and, surprisingly, that growth on an agar surface represses the cell shape phenotype of *tnr3* for at least 2 subsequent passages in broth.

## Discussion

In this study, we assembled a progenitor collection of ∼30,000 *B. theta* transposon mutants and condensed it into an ordered collection representing disruption of 2,565 genes. Our approach was relatively straightforward and should be generally applicable to most mutant libraries. Nonetheless, there were several factors that simplified the process considerably, including recruitment of personnel for relieving fatigue during repetitive tasks, ample space within the anaerobic chamber for pre-reduced supplies, a liquid handling robot to avoid human error, and sufficient 80 °C freezer space to store the progenitor collection and copies of the ordered library for backup and distribution. Moreover, barcoding of the transposons and pooled sequencing improved the efficiency and accuracy of mutant location assignments in the plates (30).

Statistical analysis of the progenitor collection revealed factors that limited the assembly efficiency of sorting and final coverage of the condensed collection. Biases in strain abundance across the pooled library resulted in the repeated isolation of some barcodes. Furthermore, we ignored barcodes associated with more than one insertion site, which reduced the number of strains that were available for re-arraying into the condensed collection. We expect that an important step toward minimizing biases in strain abundance will be to limit outgrowth of the pooled library before using it as an initial pool during progenitor collection assembly. In addition, alternative transformation strategies such as electroporation of *in vitro*-assembled transposomes (24) could reduce the occurrence of strains with multiple insertions. While the high diversity of pooled libraries and deep sequencing minimizes the impact of these issues on pooled fitness assays, efficient assembly of ordered collections has more stringent requirements that motivate further enhancement of transformation protocols. Further optimization of the pooling protocol and downstream analysis is also likely to improve assembly efficiency. Importantly, the effects of these factors on assembly efficiency could be predicted quantitatively from the statistics of barcodes in the pooled library (Figure 3), indicating that the necessary collection size, optimal pooling strategy, and other aspects of the assembly of ordered collections can be anticipated in the future.

We used a cell sorter to generate the progenitor collection due to its speed (∼3,000 wells per hour) and ability to isolate single cells. To ensure mostly axenic cultures, we used stringent gating parameters that were likely to exclude cells with aberrant morphologies, such as the filamented cells expected for cell division and DNA replication mutants (42, 43). Nonetheless, mutants such as BT2397 (*tnr3*) and cell division mutants with heterogenous cell morphologies (44, 45) can still be isolated using this approach, albeit perhaps at lower frequency. Alternative isolation strategies such as colony picking robots are less likely to bias ordered collections away from abnormal cell shapes, but the effects of colony picking on the parameters controlling assembly efficiency still need to be characterized. Changes to the sorting protocol, for instance relaxing gating parameters or sorting stationary-phase cells that are more homogenous in size (46), could expand the range of cell morphologies represented in the progenitor collection, and can be included as addendums to the progenitor collection to avoid biasing the primary sort. Since the gut microbiota is composed of species with diverse cell morphologies, the isolation of mutants with aberrant cell shapes will be important for understanding the role of cell shape in gut colonization, biogeography, and niche partitioning.

Intriguingly, nearly all mutants exhibited wild-type-like growth and shape phenotypes in the nutrient-rich medium BHIS, similar to screens of *E. coli* gene deletions in the nutrient-rich medium LB (34, 35). Nonetheless, condensed collections enable physiological screens that can illuminate previously uncharacterized roles of genes and pathways. We identified a mutant in BT0870 (*spt*) that exhibited both growth (Figure 4B-D) and cell shape phenotypes (Figure 5B,C). A targeted BT0870 (*spt*) deletion was previously reported that exhibited minor growth defects (39); our growth data, coupled with the observation that this strain was the sole insertion mutant in BT0870 (*spt*) in the original pooled transposon library, suggests that loss of BT0870 (*spt*) function is deleterious and that mutants may be prone to accumulation of suppressor mutations. The filamentous phenotype of some BT2397 (*tnr3*) mutant cells suggests that thiamine pyrophosphokinase plays an important role in cell shape determination in *B. theta*, as it does in fission yeast (47). Surprisingly, during liquid growth after passaging BT2397 (*tnr3*) through a colony, we observed reversion to wild-type phenotypes, indicating that growth conditions can play a major role in *B. theta* phenotypes. In the future, it will be interesting to screen the condensed collection in diverse media conditions to uncover additional phenotypes, as a previous screen of a CRISPRi library of *Bacillus subtilis* essential-gene depletions uncovered a wider distribution of phenotypes in a minimal medium compared with LB (48).

The condensed *B. theta* mutant collection we reported here should prove useful for addressing numerous phenotypes beyond growth and morphology that cannot be addressed in a pooled format, including the effect of gene disruption on other species or on metabolite production, as well as mechanistic inquiries about a mutant of interest. Indeed, mutants from our collection have already proven useful for follow-up studies to pooled transposon library screens (26) and to metabolomics measurements (29). The ordered collections should thus serve as a resource for identifying genotypes important for *B. theta*’s function within microbial communities and in the gut. Moreover, the approach we have developed for the assembly of ordered collections should be generally applicable in any organism for which diverse pooled transposon libraries can be generated, enabling genetics-based interrogation of members of the gut microbiota and the molecular mechanisms through which they impact human health.

## Methods

### Oligos and strains

The transposon-insertion mutants in the condensed collection are listed in **Dataset S1**. The genes that are only represented by multi-insertion strains, and the location of these strains in the progenitor collection, are listed in **Dataset S2**. The strains and oligos used in this study are listed in **Table S1**. Details of the transposon design, transformation, and the original Tn-seq mapping experiment were published previously (26).

### Preparation of the pooled library for sorting of the progenitor collection using flow cytometry

A barcoded transposon mutant library of *B. theta* VPI-5482 mutants was sorted using a fluorescence activated cell sorting (FACS) machine, as described previously (30) with a few modifications. Cells were sorted into 262 96-well plates in three batches of 60, 100, and 102 plates; the same protocol was used for each batch. These 262 plates were combined with 40 96-well plates that were generated in a pilot experiment (30), leading to the construction of the final progenitor collection of 302 96-well plates. All media and plasticware were pre-reduced in an anaerobic chamber (Coy Laboratories) for 3 days before use to eliminate any residual oxygen.

### Addition of medium to the 96-well plates before sorting

333 microliters of pre-reduced and filter-sterilized BHIS (with no added cysteine) were added to 2-mL 96-deep well plates (Greiner Bio-One, Cat. #1986-2110) using a semi-automated BenchSmart pipettor (Mettler-Toledo) installed in an anaerobic chamber. Each 96-deep well plate with added medium was sealed with a foil seal (Nunc^TM^ Sealing Tapes, Fisher Scientific, Cat. # 232698) and stored anaerobically for 16-24 h at 37 °C before use in the next step of the procedure.

### Outgrowth of the randomly barcoded transposon mutant library

A cryostock of pooled library (1 mL of OD600=1) was thawed in an anaerobic chamber and added to 100 mL of BHIS (26) in a 250-mL Erlenmeyer flask that had been pre-warmed to 37 °C, and the culture was incubated overnight (12–18 h) at 37 °C without agitation. The next day, ∼3 h before cultures were transported to the FACS machine, this overnight culture was diluted to an OD600∼0.1 in 4 mL of BHIS pre-warmed to 37 °C in 4 independent samples. At the same time, BHIS was pre-aliquoted into an additional set of tubes (2 mL per tube) and kept at 37 °C.

### Sorting the pooled library to create the progenitor collection

Immediately before being transported to the FACS machine, log-phase cultures of the transposon library were diluted to an OD600 of 0.01-0.05 in pre-warmed BHIS with 10 mM cysteine in a FACS tube (Falcon® high-clarity polypropylene round bottom test tubes, Corning, Cat. #352063) and mixed thoroughly in the anaerobic chamber. An initial culture in a FACS tube was brought out of the chamber and loaded onto the FACS machine to calibrate the instrument and define a gate based on forward and side scatter. After calibrating the machine, a fresh FACS tube of culture and a set of pre-reduced 96-deep well plates were brought to the FACS machine and the fresh culture was used to sort single cells. To preserve the anaerobic environment of the culture, the FACS tube was sealed in an airtight container inside the anaerobic chamber for transportation to the FACS machine, and the FACS tube was disturbed as little as possible after being exposed to oxygen. After single cells were sorted into individual wells of a 96-deep well plate, the deep well plate was lightly resealed with a gas-permeable seal (Excel Scientific Inc., Cat. #BS-25). Batches of 15 96-deep well plates with sorted cells were returned to the anaerobic chamber and new sets of deep well plates with fresh media were transported to the FACS facility as needed to keep the FACS machine in continual operation. Once sorted plates were returned to the anaerobic chamber, the gas-permeable seal was fully sealed using a rubber brayer roller, taking care to seal the edges of the plate to prevent evaporation. The sorted plates were then returned to the 37 °C incubator in the anaerobic chamber. The FACS tube of culture was replaced with a fresh culture every 30 plates. Cultures of the transposon library were kept in log-phase via dilution in the 37 °C incubator throughout the course of the sorting experiment to maintain a supply of log-phase cells for sorting.

### Aliquoting copies of cryo-stocks of the progenitor collection

The sorted cells were allowed to grow into monocultures over 2 days. Glycerol was added to the cultures, and the glycerol stock was aliquoted into two copies of the library using a semi-automated 96-well BenchSmart pipetting robot inside the anaerobic chamber. Eighty microliters of 50% glycerol were added to each well (final concentration 15% glycerol) and mixed by pipetting up and down. Eighty microliters of the glycerol stock were then aliquoted into two V-bottom 96-well plates (Greiner Bio-One, Cat. #651161). The cryo-stock plates were sealed with a foil seal and stored at -80 °C. The remainder of the glycerol stock inside the 96-deep well plate was also stored for subsequent pooling steps of the protocol.

### Re-arraying the progenitor collection

At the beginning of the experiment, erythromycin (Sigma, Cat. #E5389-5G) was added at a final concentration of 10 µg/mL to a bottle of filter-sterilized BHIS (Becton Dickinson, Cat. #237500) without cysteine that had been left in the anaerobic chamber for 48-72 h to reduce. This selective medium was aliquoted into 8 96-deep well plates (Celltreat, Cat. #229574) using a BenchSmart with a 1000-µL head (Rainin, Cat. #BST- 96-1000), sterile filter tips (Rainin, Cat. #30296782), and a 300-mL reservoir (Integra, Cat. #6328). The 96-deep well plates were sealed with a plastic film (Excel Scientific, Cat. #STR-SEAL-PLT), transferred to a second anaerobic chamber using airtight plastic boxes, and stored in a 37 °C incubator inside the anaerobic chamber until inoculation.

Over the course of a day, sections of the progenitor collection were re-arrayed anaerobically using an Eppendorf EpMotion 5073 (Eppendorf, Germany) running EpBlue v. 40.5.3.10 and housed inside an anaerobic chamber. Table S2 contains the settings used for the EpMotion.

Before transfer to the anaerobic chamber, the 96-well V-bottom plates containing glycerol stocks of the progenitor collection were removed from the -80 °C freezer in batches and stored on a bed of dry ice. The lids of each plate were removed, ice was cleaned off, and the lids were sterilized with 70% (v/v) ethanol while the sealed plates were centrifuged in a microplate centrifuge (Fisherbrand, Cat. #14-955-300) for 30 s. After centrifugation, the foil seal was removed from each plate, taking care not to jostle the plate. The clean and sterilized lid was then returned to each plate, and the lidded plates were returned to the bed of dry ice. When the entire batch of plates had been processed in this manner, the plates were transferred into the anaerobic chamber and stored in a safe location on the benchtop.

Next, we inoculated the 96-deep well plates of selective medium with 40 µL of glycerol stock from selected wells of the progenitor collection. Three hundred microliter PCR-clean filter tips (Eppendorf, Cat. #0030 014.472) were used in combination with a single channel 300-µL adaptor (Eppendorf) to transfer glycerol stocks. The pipetting pattern (the set of instructions connecting position in the progenitor collection to position in the condensed collection) was imported into EPBlue as a .csv file after being generated using a Matlab script.

After inoculation, 96-well plates from the progenitor collection were re-sealed with a foil seal (Thermo Scientific, Cat. #232699) and transferred back to the -80 °C freezer. The 96-deep well plates with freshly inoculated cultures comprising the condensed collection were sealed with a gas-permeable seal (Excel Scientific Inc., Cat. #BS-25) and stored in a 37 °C incubator in the anaerobic chamber to recover for 36-48 h and used to inoculate growth curves and to aliquot copies for cryostorage.

### Growth curve inoculation

Approximately 48 h post-inoculation, deep well cultures were used to inoculate fresh medium for growth curve measurements and then aliquoted into glycerol stocks as copies of the final condensed collection (glycerol stock storage described below).

First, 198 µL of BHIS without cysteine and without erythromycin was aliquoted across 16 96-well flat-bottom plates (Greiner Bio-One, Cat. #655180) using a BenchSmart with a P1000 head and sterile filter tips. The plates were transferred to the anaerobic chamber along with the previously generated cultures using a airtight sealed container.

The cultures were then used to inoculate fresh medium for growth curves using an EpMotion 5073. Fifty microliter PCR-clean filter tips (Eppendorf, Cat. #0030 014.430) were used in combination with an 8-channel 50-µL volume adaptor (Eppendorf). Each deep-well culture was used to inoculate 2 96-well flat-bottom plates as replicates for the growth curve measurements. Two microliters of culture were transferred without mixing at the source and with 1 mixing step of 40 µL at the target. The same tips were used, and the source was revisited once, to inoculate a replicate target plate. Table S3 contains the machine settings used for this protocol. To avoid transferring liquid from the intentionally blank wells on each plate, we removed the tips from positions A1, B1, and the other blank well on the plate (Dataset S1).

For some of the flat-bottom growth curve plates, 2 µL of a culture of wild-type *B. theta* VPI-5482 grown in BHIS without cysteine for 36-48 h were used to inoculate position A1 as a positive control. All flat-bottom 96-well plates were sealed with modified sterile plastic seals, cut to not extend over the edges of the plates, and assembled in a plate stacker (BIOSTACK3WR, Biotek Instruments Inc., Cat. #05404-0998) associated with a Synergy H1 microplate reader (Biotek Instruments Inc.) running Gen 5 v. 3.08.01. The plate stacker and the front of the microplate reader were enclosed in a custom-fabricated box along with a thermal control unit (AirTherm SMT, World Precision Instruments) to ensure a constant temperature of 37 °C during growth curve measurements. The plate stacker constantly read plates and one complete run through all plates required 30-42 min depending on the number of plates. The plate reader settings were as follows: 37 °C, 10 s of shaking at 282 cycles per min with a double orbital pattern before reading optical density at 600 nm. After approximately 48 h of growth, the plates were removed from the plate stacker and used for single cell imaging.

### Storing glycerol stocks

After being used to inoculate flat-bottom plates for growth curves, the deep-well cultures were sealed with a plastic film (Excel Scientific, Cat. #STR-SEAL-PLT) and transferred back to an anaerobic chamber using an airtight box. The BenchSmart 96-well pipetting robot was used along with a P1000 head and a 300-mL reservoir to transfer 353 µL of a sterile solution of 50% glycerol (Fisher, Cat. #G33-4) and 50 mM cysteine (Millipore, Cat. #243005-100GM) that had been pre-reduced inside the chamber for >48 h. After mixing the cultures with glycerol by pipetting up and down twice, the glycerol stocks were distributed in 80-µL aliquots into 96-well V-bottom plates covered temporarily with a sterile lid (Greiner, Cat. #656161) as copies of the final condensed library. Aliquoted library copies were sealed with a foil seal, the sterile lid was placed over the seal, and transferred to a -80 °C freezer for long-term storage.

### Pooling strategy

Wells in the progenitor collection were pooled according to a plate-well strategy that requires *N*+96 pools, where *N* is the number of 96-well plates. Pooling essentially followed the procedure described previously (30). Individual wells were first pooled, either pooling the same well from all plates (e.g. A1 from progenitor collection plates 41–302) or pooling all wells from a single plate (e.g. A1–H12 from plate 41). In previous efforts (30), the first pool set (same well, different plates) was pooled further to create 8 and 12 row and column pools, respectively. In this work, the set of pools from the 96 well positions was sequenced directly, along with the 262 plate pools. As described previously (30), a single pool was made from every well of the progenitor collection extension (plates 41–302) for use as input to RB-TnSeq.

With the plate-well pooling strategy used here, the location of a barcode isolated *n* times in the collection will be narrowed down to *n*^2^ possible wells. For example, a barcode isolated at position G1 of plate 1 (P1-G1) and position H2 of plate 2 (P2-H2) of the progenitor collection will be sequenced in pools P1, P2, G1, and H2. The four possible locations that are consistent with these results are P1-G1, P1-H2, P2-G1, and P2-H2. Two of the possible locations in this example are the true locations, while the remaining two are artifacts of the pooling and decoding process.

When a barcode did not have a definite location, we used a probabilistic strategy to predict the likelihood of a particular configuration of wells, as described previously (30, 32). Critically, this algorithm depends on systematic differences in the contribution of each well in the pool to the total number of sequencing reads (32). While BarSeq is particularly effective compared to other methods at providing a quantitative and accurate estimate of the relative abundance of a barcode in the pool (30), plate-well pools carrying similar relative abundance of the barcode in question are poor in information and hence the resulting predictions are low in confidence. Here, we used a heuristic cutoff of 0.85 for considering a predicted configuration of locations to be high confidence.

### Sequencing the progenitor collection

DNA from plate-well pools of the progenitor collection was isolated with a DNeasy 96 Blood &Tissue Kit (Qiagen, Cat. #69582) according to manufacturer’s instructions. BarSeq was performed on individual pools with indexed primers (Table S1), as described previously (30), and sequenced on a MiSeq (Illumina, SY-410-1003) using MiSeq Reagent Kit v3 (150-cycle) (Illumina, MS-102-3001). DNA was isolated from the complete pool at the same time and used as input for an RB-TnSeq protocol, as detailed previously (30) (Table S1). The RB-TnSeq library was sequenced on a MiSeq using MiSeq Reagent Kit v3 (150-cycle).

### Decoding the progenitor collection

The process of locating barcodes within the progenitor collection was performed essentially as previously described for the initial 40-plate collection (30), with small modifications to account for the change in pooling strategy from row-column-plate to plate-well. Briefly, we used the BarSeq results from individual pools to locate barcodes within the collection. Barcodes with definite solutions (isolated once in the collection) were identified first, then statistics on the distribution of abundance of these barcodes were used to inform the likelihood of solutions for the location of barcodes without definite solutions (isolated more than once in the collection).

Simultaneously, we incorporated the RB-TnSeq data of the progenitor collection into the larger RB-TnSeq dataset from the initial pooled library. The higher depth of sequencing from RB-TnSeq on the progenitor collection allowed us to map more barcodes to the genome and provided higher sensitivity for the detection of multiple insertion sites associated with the same barcode. Once a barcode was located in the collection, a lookup table connecting barcodes to insertion sites was used to determine its utility as a mutant strain for the condensed collection.

The detailed algorithm for determining the mapping status of a barcode (e.g. single insertion vs. multiple/ambiguous insertion) can be found in the code published previously (30). Importantly, we only considered insertion locations in the genome for which the number of reads was >25% that of the most abundant insertion location for the same barcode. While the same barcode mapping to multiple locations in the genome in a pooled library could arise from multiple causes (such as the chance occurrence of the same barcode in multiple strains), RB-TnSeq of the pooled progenitor collection indicated that the majority of barcodes that mapped to multiple sites were isolated only once, consistent with previous results (30). If the insertions of these ambiguous barcodes were found in separate strains, this scenario would require the repeated sorting of two or more cells with the same barcode into the same well, but the occurrence of wells in the collection with different barcodes was rare (2%). Therefore, in this study we treated barcodes associated with more than one insertion site as multiple-insertion strains and excluded them from re-arraying into the condensed collection.

### Modeling assembly of the progenitor collection

To quantify barcode abundance in the initial pool, a 1 mL-aliquot of the same initial pool as the one used for sorting the additional 262-plates was inoculated into BHIS and recovered overnight at 37 °C. Six 1-mL aliquots of this overnight culture were pelleted, DNA was extracted, and BarSeq was performed as above (see **Sequencing the progenitor collection**). This protocol is the same as the one used to generate the *t0* samples that serve as controls for pooled fitness assays, and we expect that any *t0* sequencing data will be useful for modeling collection assembly as long as it comes from the same initial pool as the culture used for sorting. Barcode abundances were summed from these *t0* samples and the relative abundance across the summed pools was calculated and used as a probabilistic weight for random sampling during simulations. Before drawing from the pool, we filtered out barcodes in the *t0* samples that were not associated with any insertion location (unmapped).

To simulate the assembly of a collection, *b* barcodes were randomly drawn with replacement from this initial set, insertion sites were located, and genome coverage was determined. We required that an insertion be found between the first 5% and last 10% of a gene to consider that gene disrupted. We simulated a range of collection sizes (*b*) and performed 250 simulations for each collection size. To account for assembly efficiency (*K*), we scaled the collection size in the simulated coverage curves by *K*^-1^. To assess the impact of strain abundance bias, we simulated collection assembly from an initial pool in which the weights were set to be equal. To assess the impact of assembly bias, we set *K*=1.

The value of *K* was estimated by quantifying three parameters from the statistics of the 262-plate library: *b*/*w*, the number of single barcodes per well; *f*mapped, the fraction of barcodes mapped in RB-TnSeq; and *f*single, the fraction of barcodes associated with a single site. These parameters were chosen because they represent the filtering steps used in this work to determine whether a barcode was useful for inclusion in the final condensed collection. We expect that *K* can be estimated for any protocol, as long as the fraction of wells with a useful barcode is accurately quantified. We computed the best-fit value of *K* (*K*fit) using a custom Matlab script that minimized the sum of squares of the difference between true(*w*), the true saturation curve of size *w* wells, and sim(*K*fit *b*), the simulation of size *b* barcodes scaled by the fitting parameter *K*fit . The experimentally estimated value of *K* was used as the initial condition, although the final solution of fitting was robust to changes in the initial condition.

### Measurement of population growth metrics

Maximum growth rate was calculated as the largest slope of ln(OD) with respect to time (calculated from a linear regression of a sliding window of 11 time points) using custom Matlab (Mathworks, Natick, MA, USA) code.

### Single-cell imaging

Stationary-phase cells were diluted 1:10 into 0.85X PBS and then taken from 96-well plates and placed on 1% agarose pads with 0.85X PBS to control for osmolality. Phase-contrast images were acquired with a Ti-E inverted microscope (Nikon Instruments) using a 100X (NA 1.40) oil immersion objective and a Neo 5.5 sCMOS camera (Andor Technology). Images were acquired using µManager v.1.4 (49). High-throughput imaging was accomplished using SLIP, as described previously (38). Including sample preparation and calibration, SLIP enables acquisition of 49 images per well of a 96-well plate in ∼30 min. Since replicate growth curves appeared similar across the entire library (Figure S2), we imaged one replicate culture for each gene.

### Morphological analyses

The MATLAB image processing code *Morphometrics* (35) was used to segment cells from phase-contrast or fluorescence microscopy images. A local coordinate system was generated for each cell outline using a method adapted from *MicrobeTracker* (50). Cell widths were calculated by averaging the distances between contour points perpendicular to the cell midline, excluding contour points within the poles and sites of septation. Cell length was calculated as the length of the midline from pole to pole. Cell surface area was estimated from the local meshing.

### Growth curve measurements from single colonies

To isolate single colonies, we aerobically struck glycerol stocks onto BHIS 1.5% agar plates and then transferred the plates to an anaerobic chamber and incubated them at 37 °C for 48 h. We performed all further steps in an anaerobic chamber. We inoculated single colonies into a pre-reduced and pre-blanked flat bottom plate (Greiner Bio-One, Cat. #655180) with 200 µL of pre-reduced BHIS and into a 96-deep well plate (Greiner Bio-One, Cat. #1986-2110) with 500 µL of pre-reduced BHIS using and incubated the cultures for 48 h. The flat-bottomed plate was used to measure the outgrowth from the colony and the deep well plate was incubated without shaking in a 37 °C incubator. Since BT0870 colonies were visible but much smaller than wild type, we combined 5-6 colonies of the BT0870 mutant into one culture to approximately normalize the inoculum density. 2 µL of deep well cultures were used to inoculate a pre-reduced and pre- blanked flat bottom plate with 200 µL of pre-reduced BHIS. For both outgrowth from a colony and from the 48 h cultures measurements, we applied an optical seal (Excel Scientific, Cat. #STR-SEAL-PLT) and measured OD600 with a Biotek Epoch plate reader with the following settings: temperature 37 °C, reading of OD600 every 5 min with continual orbital shaking (3 mm, 282 cycles per min) between reads. We subtracted well-specific blank values before plotting the growth curves (51).

## Supporting information

Dataset S1 - Strains in condensed collection

Dataset S2 - strains with multiple insertions

## Acknowledgements

We thank Dr. Paula Welander for critical insights and members of the Huang lab for helpful discussions. This work was funded by NIH RM1 Award GM135102 (to A.D. and K.C.H.) and R01 AI147023 (to K.C.H.). K.C.H. is a Chan Zuckerberg Biohub Investigator.

## Supplementary Figures

**Figure S1:**
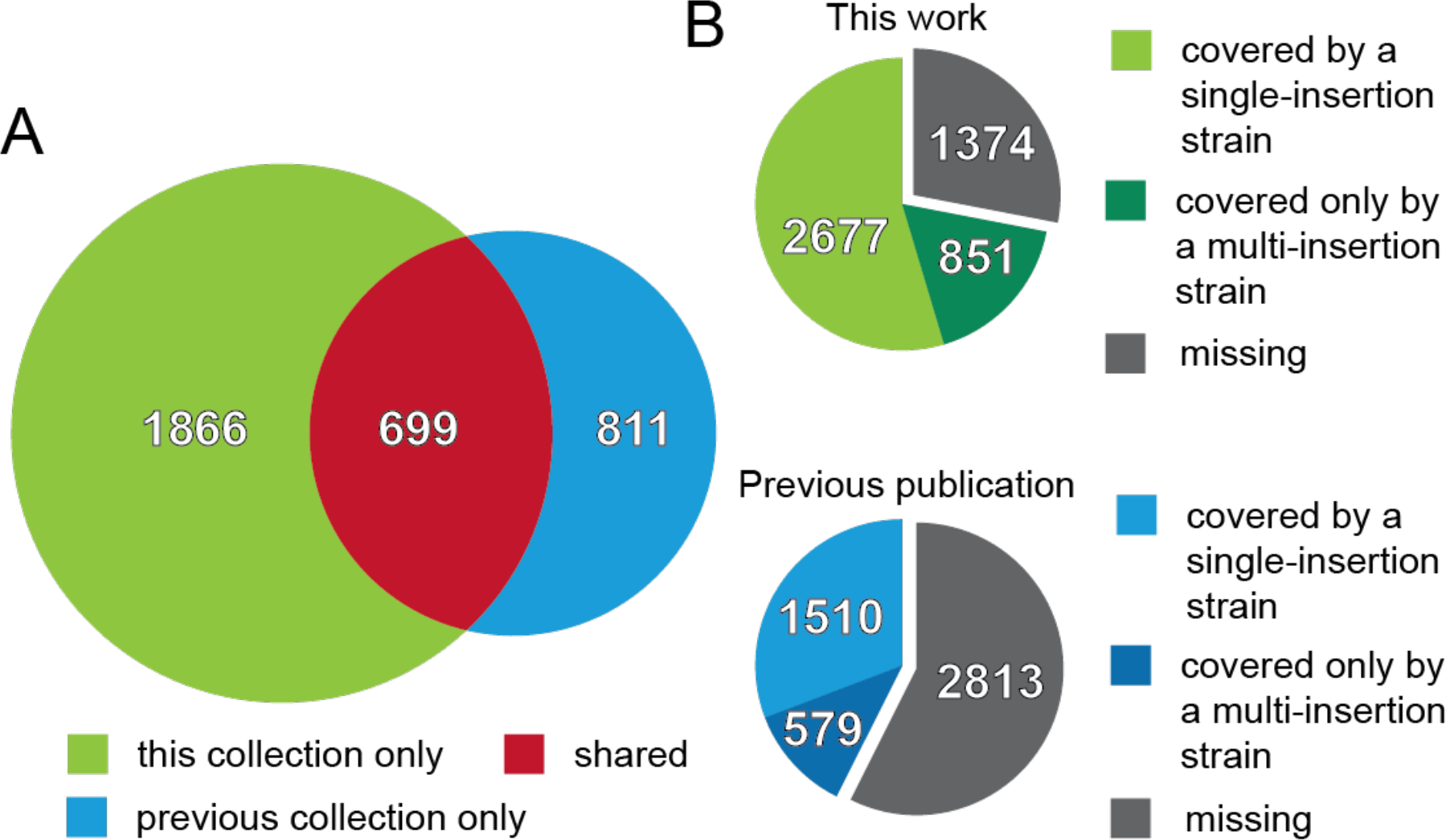
The progenitor collection expands coverage of the *B. theta* genome. A) The genes covered by single insertions in the progenitor collection overlap to some degree with a previously published (23) ordered collection of *B. theta* VPI- 5482 transposon mutants. The Venn diagram shows the number of genes that overlap between the two datasets (699) and the number that are unique to the progenitor collection (1,866) and the previously published collection (811). Information on insertion locations in the previous collection was extracted from published materials and reanalyzed using the same criteria for coverage as the collection in this work. B) The progenitor collection expands coverage of the *B. theta* genome. The number of genes represented by transposon mutant strains is shown for the progenitor collection (top) and the previously reported collection (bottom). As in (A), information on insertion location was extracted from the previous publication and reanalyzed. Even when only considering strains with single insertions, the condensed collection expands coverage by 1,167 genes.

**Figure S2:**
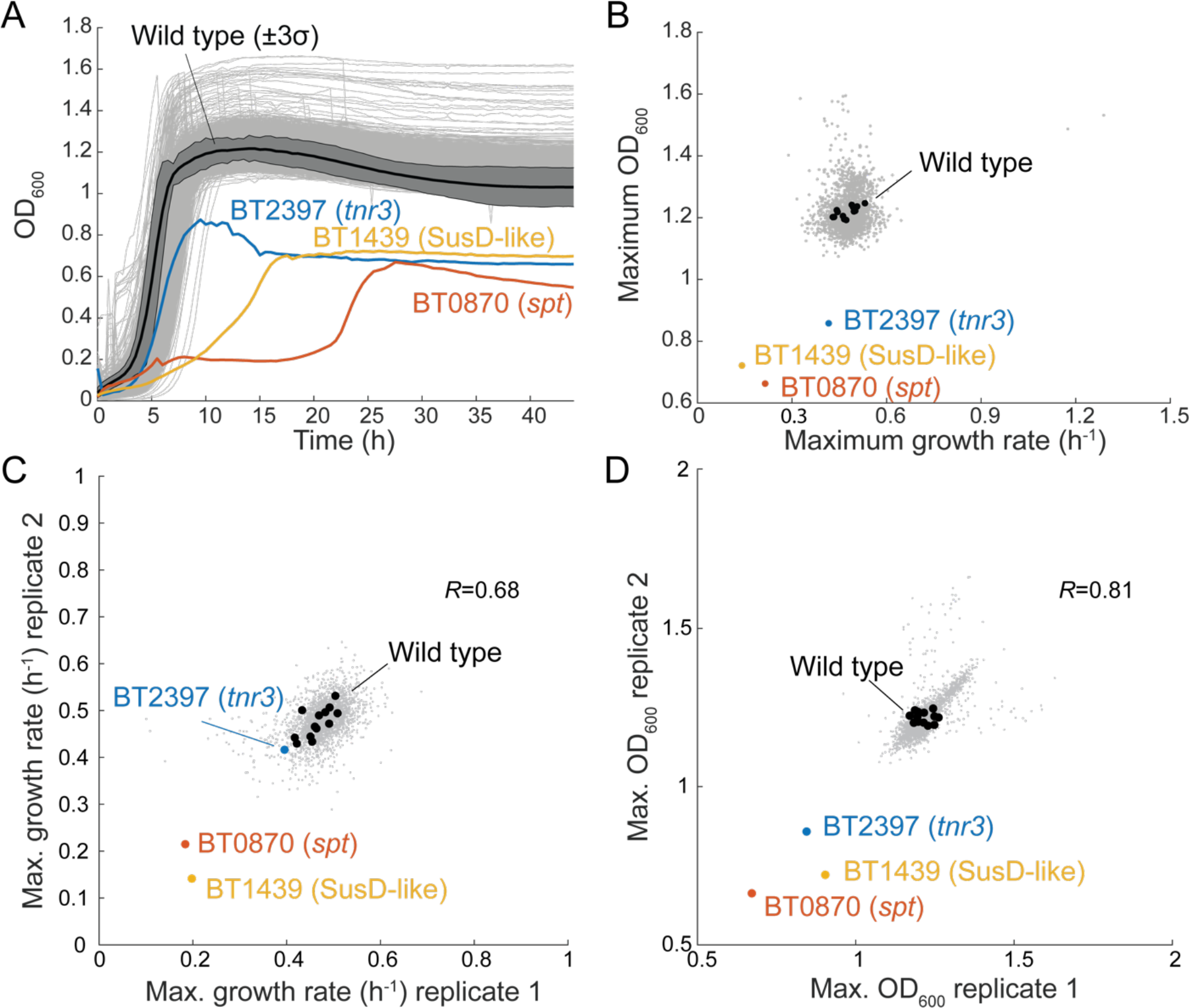
Technical replicates of growth measurements from the condensed collection consistently highlight the same set of mutants with growth defects. A) A technical replicate of growth curves of the condensed collection led to identification of the same set of mutants with growth defects. Similar growth defects were observed for BT2397 (*tnr3*, blue), BT0870 (*spt*, red), and BT1439 (SusD-like, yellow) mutants as in the first replicate (Figure 4B). The majority of strains grew similarly to wild-type controls (black, with shaded dark gray region representing 3 standard deviations). B) Maximum growth rate and maximum OD600 extracted from the technical replicate growth curves in (A) highlight the growth defects of BT2397 (*tnr3,* blue), BT0870 (*spt,* red), and BT1439 (SusD-like, yellow) mutants. C) Maximum growth rate was consistent between technical replicates of the growth curves. Pearson’s correlation coefficient *R*=0.68. Black circles are wild-type controls, which exhibit similar spread as the condensed collection. D) Maximum OD600 was reasonably consistent between technical replicates of the growth curves. Growth curves with OD600 much higher than wild type were generally not reproducible between replicates. Pearson’s correlation coefficient *R*=0.81. Black circles are wild-type controls.

**Figure S3:**
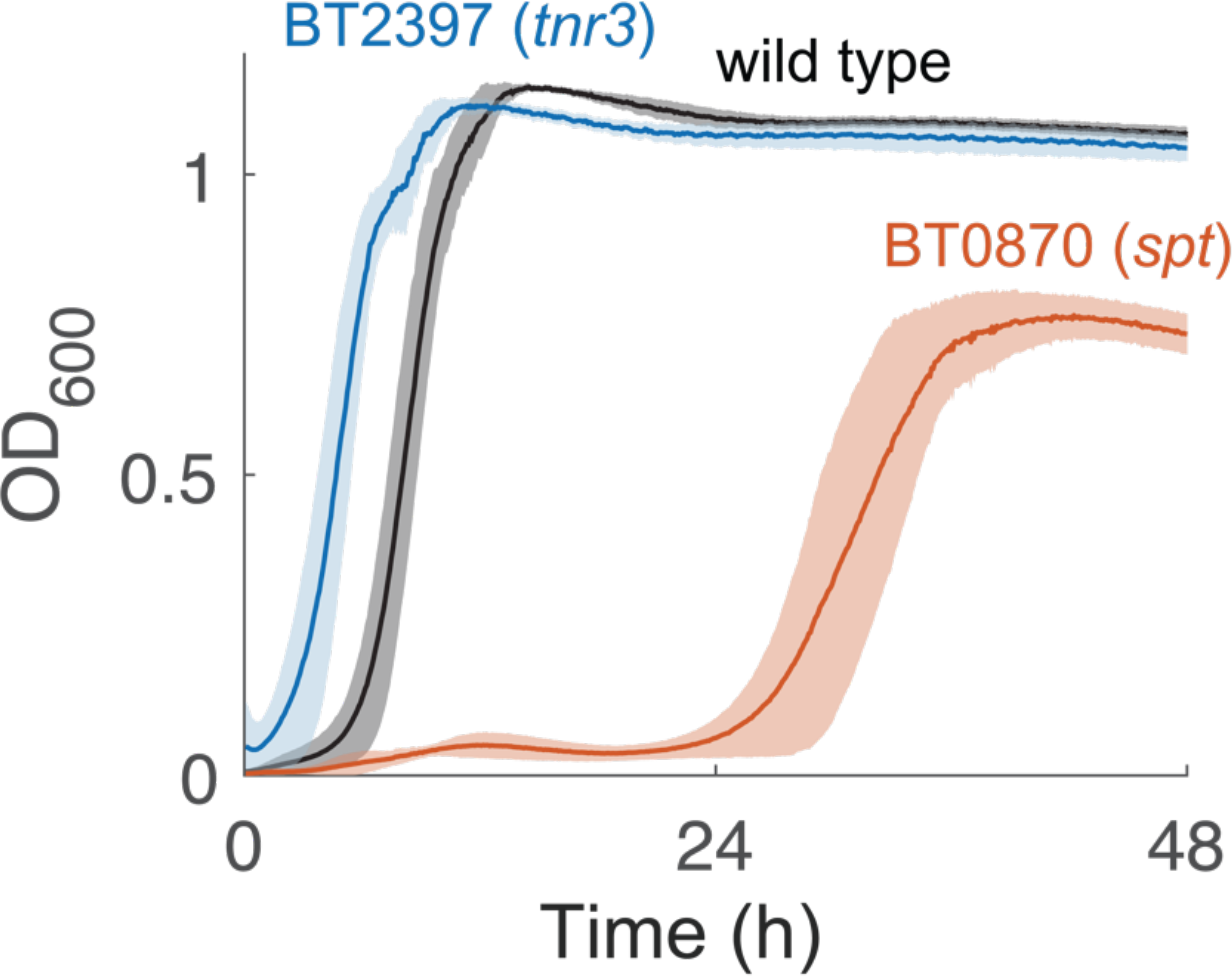
The BT0870 (*spt*) mutant exhibits growth defects during outgrowth from a colony. BT2937 (*tnr3*, blue) and BT0870 (*spt*, red) cultures inoculated directly from colonies into liquid BHIS displayed qualitatively similar growth curves to cultures inoculated from a liquid passage after colony growth (Fig. 4D). Wild type is shown in black. Shaded regions represent 1 standard deviation for *n*=6 biological replicates.

**Figure S4:**
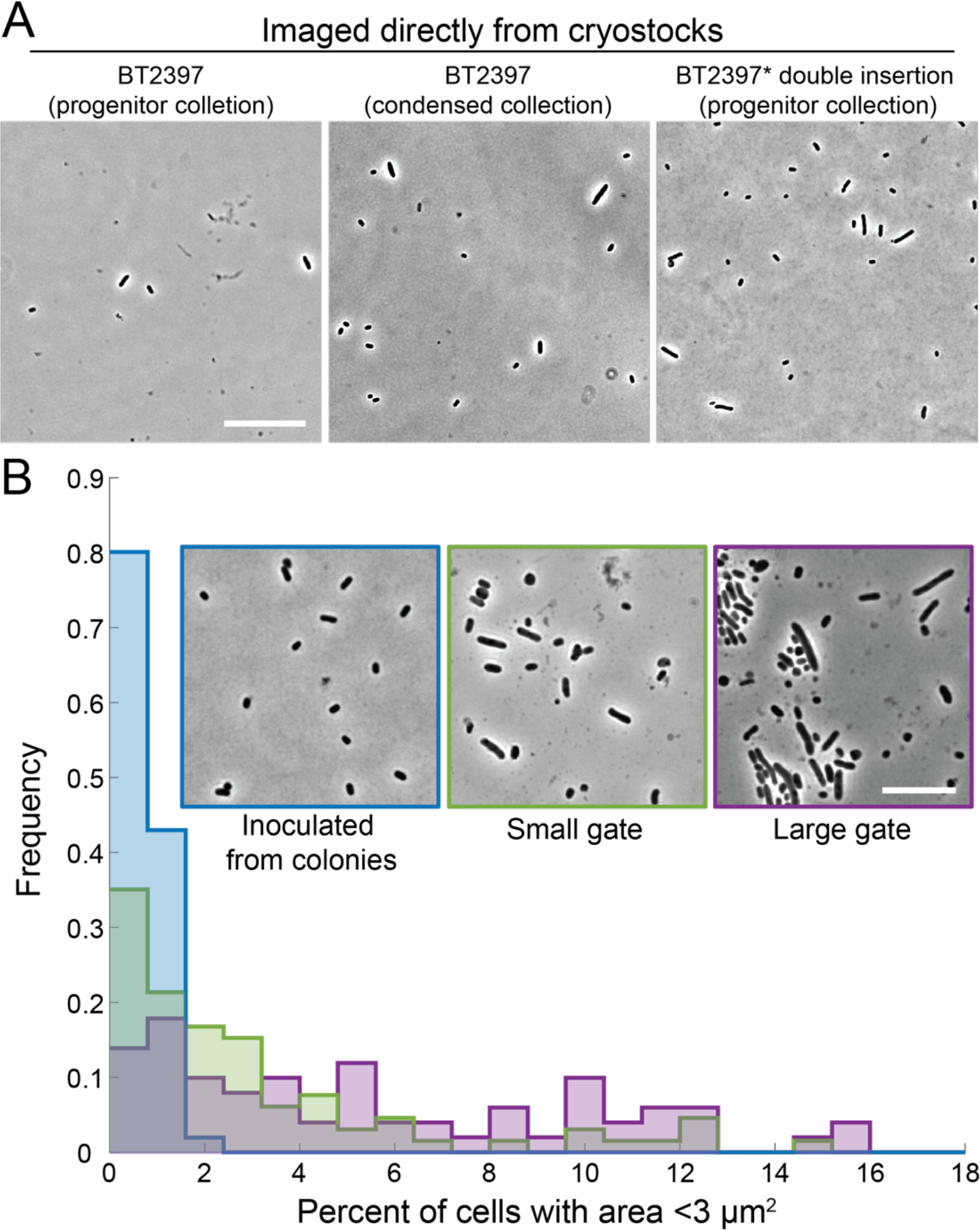
The elongated cell phenotype of the BT2937 (*tnr3*) mutant is specific to passaging through liquid. A) Independent strains with insertions in BT2397 (*tnr3*) exhibited elongated cells, including the BT2397 (*tnr3*) cryostock from the condensed collection and two independent strains from the progenitor collection (one was the single-insertion BT2397 (*tnr3*) mutant that was propagated for the condensed collection and the other has a barcode associated with insertions in BT2937 and BT2343). Cells were spotted directly onto agarose pads after dilution from the cryostock and imaged aerobically without growth. Bar: 20 µm. B) Cultures inoculated by sorting a single cell into liquid BHIS and passaged twice exhibited an increased fraction of elongated cells, while cultures inoculated from a colony and passaged twice in liquid BHIS (blue) before imaging displayed uniform, approximately wild-type shapes. Sorting was performed with either a gate to select for small, approximately wild-type shaped cells (green) or a gate to select for larger cells (purple), and cells were passaged in liquid BHIS twice before imaging; in both cases, a substantial fraction of cultures contained >2% of cells with area >3 µm^2^, unlike cultures inoculated from a colony. Representative images are shown in the inset. Bar: 10 µm. *n*=64, 83, and 63 cultures for passaging through a colony, inoculated with a small cell, or inoculated with a large cell, respectively. *n*>300 cells were segmented per culture.

## Supplementary Tables

**Table S1:**
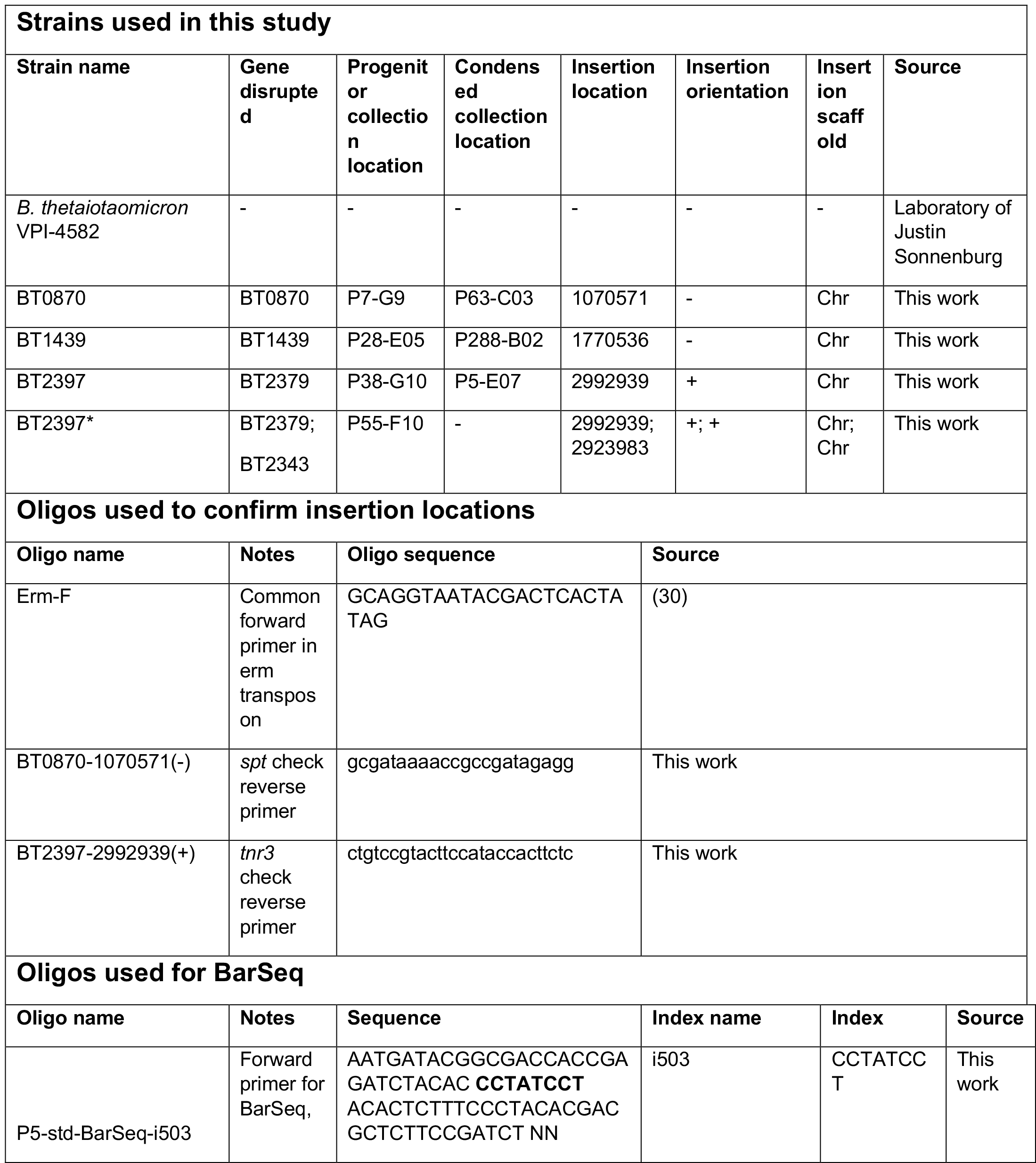
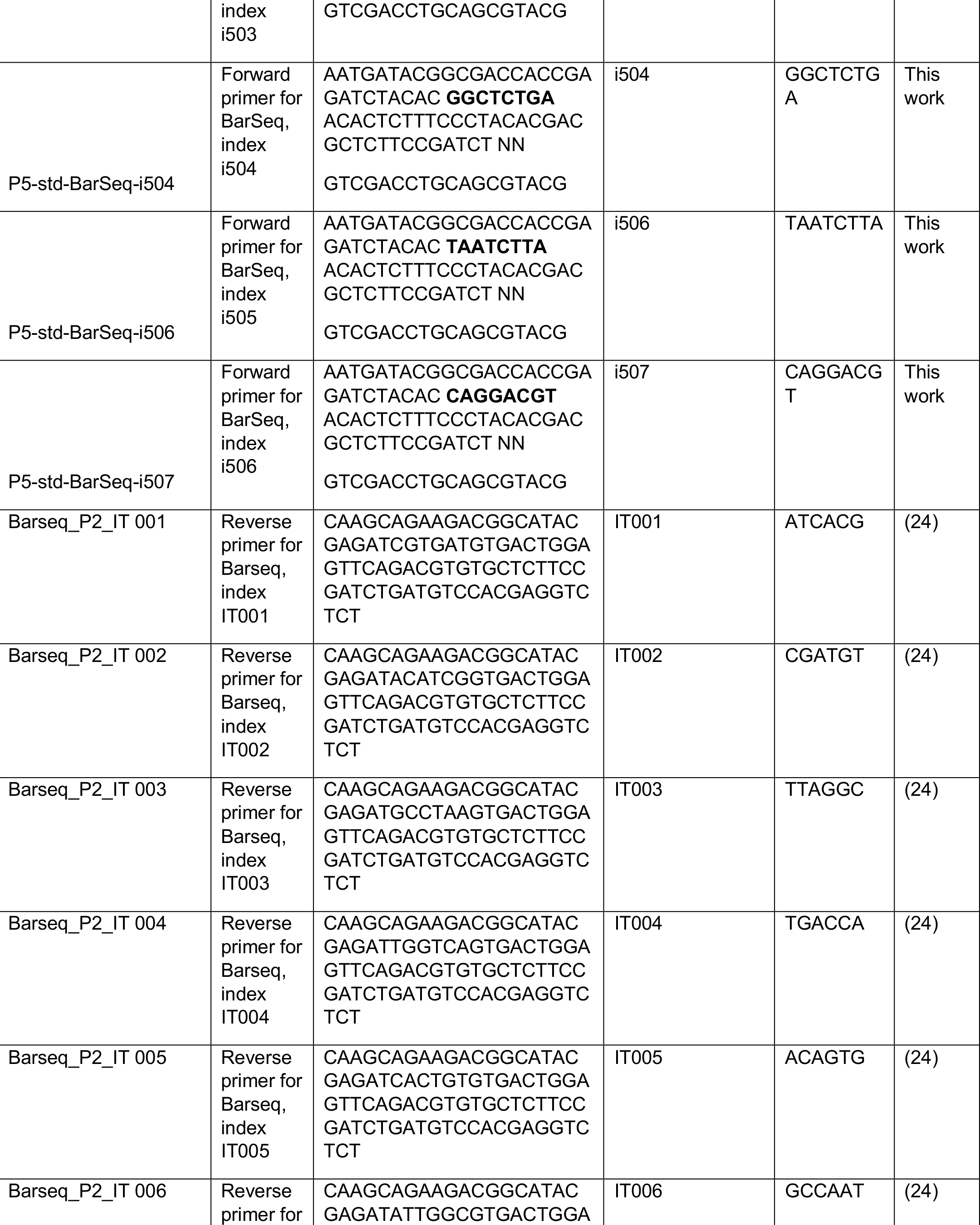
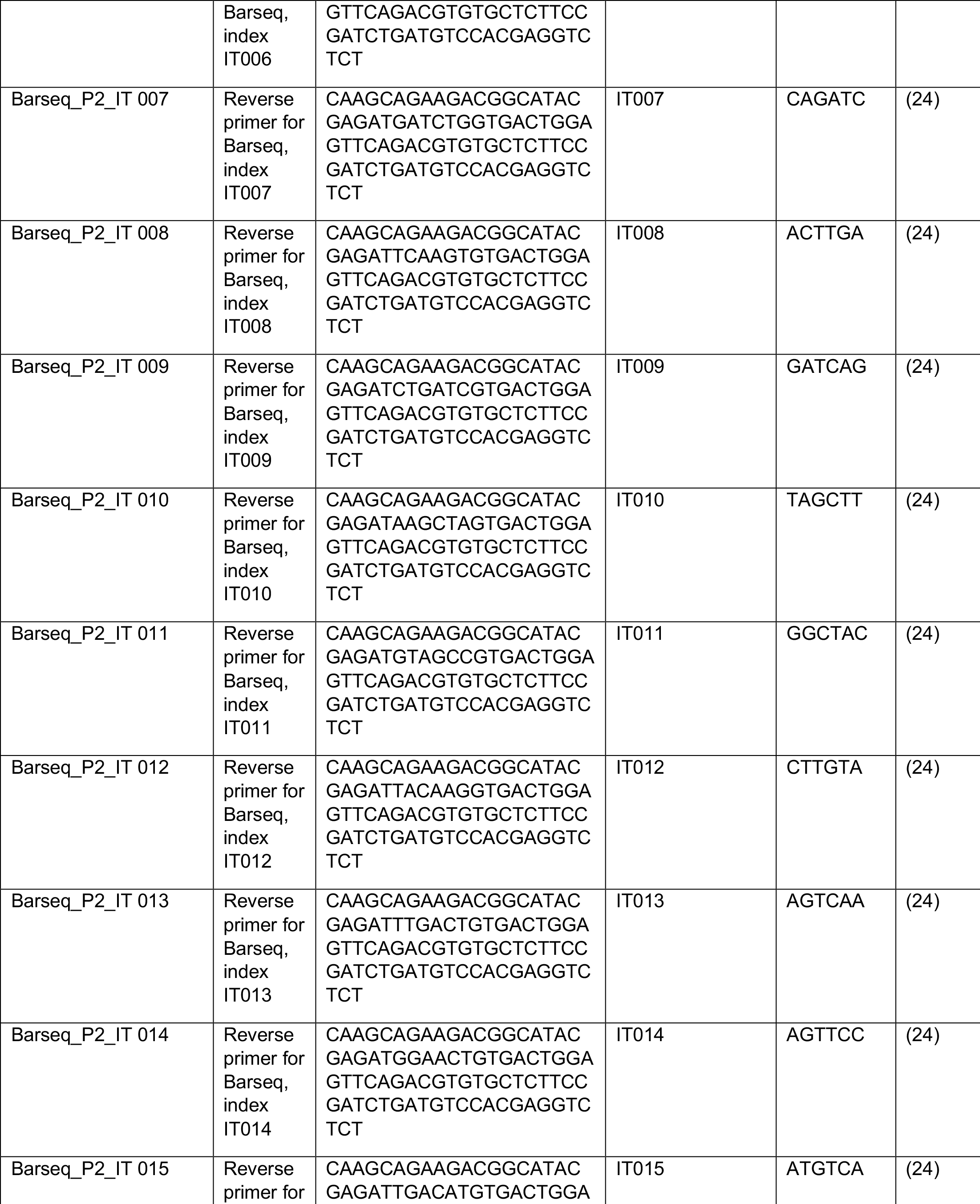
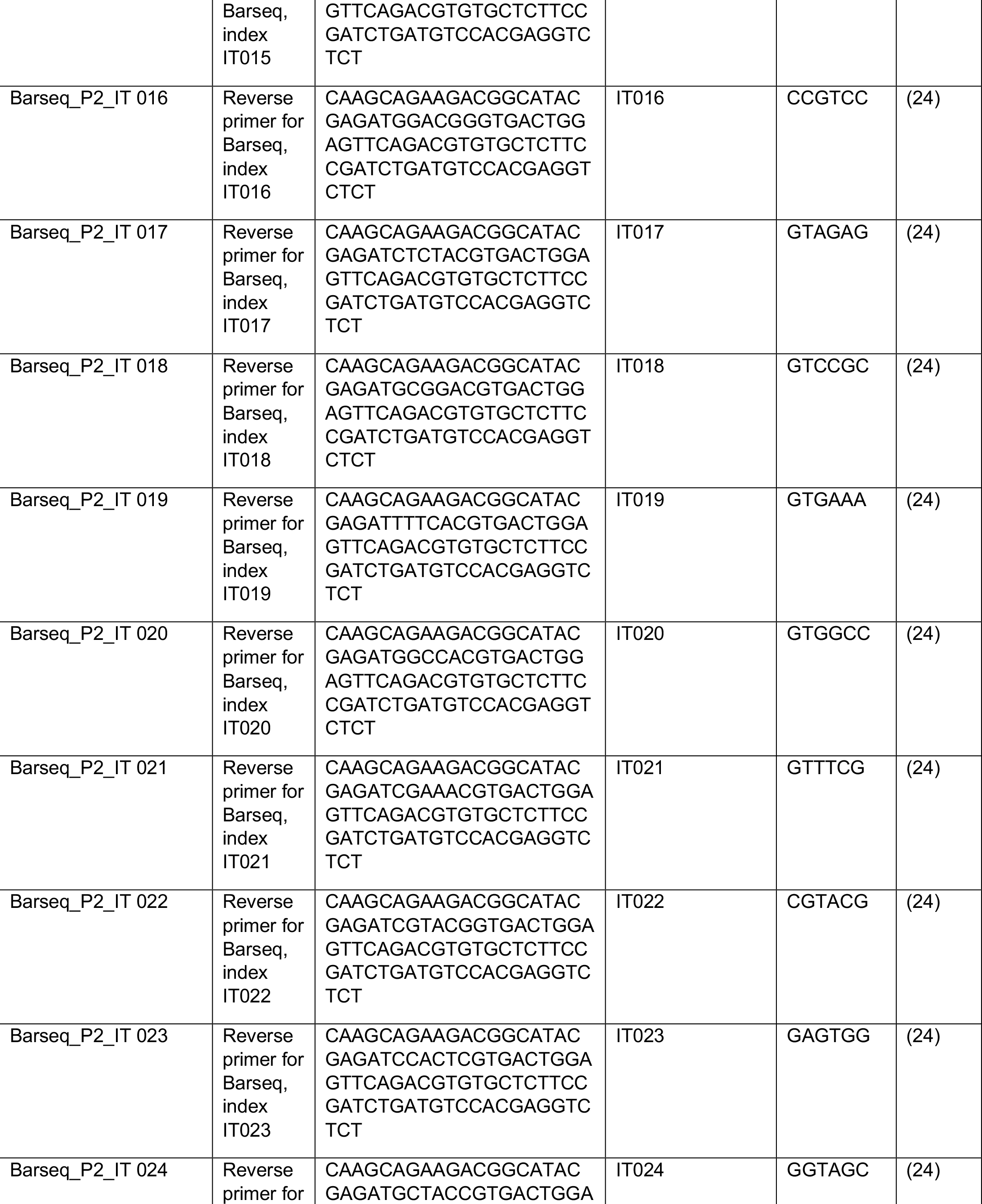
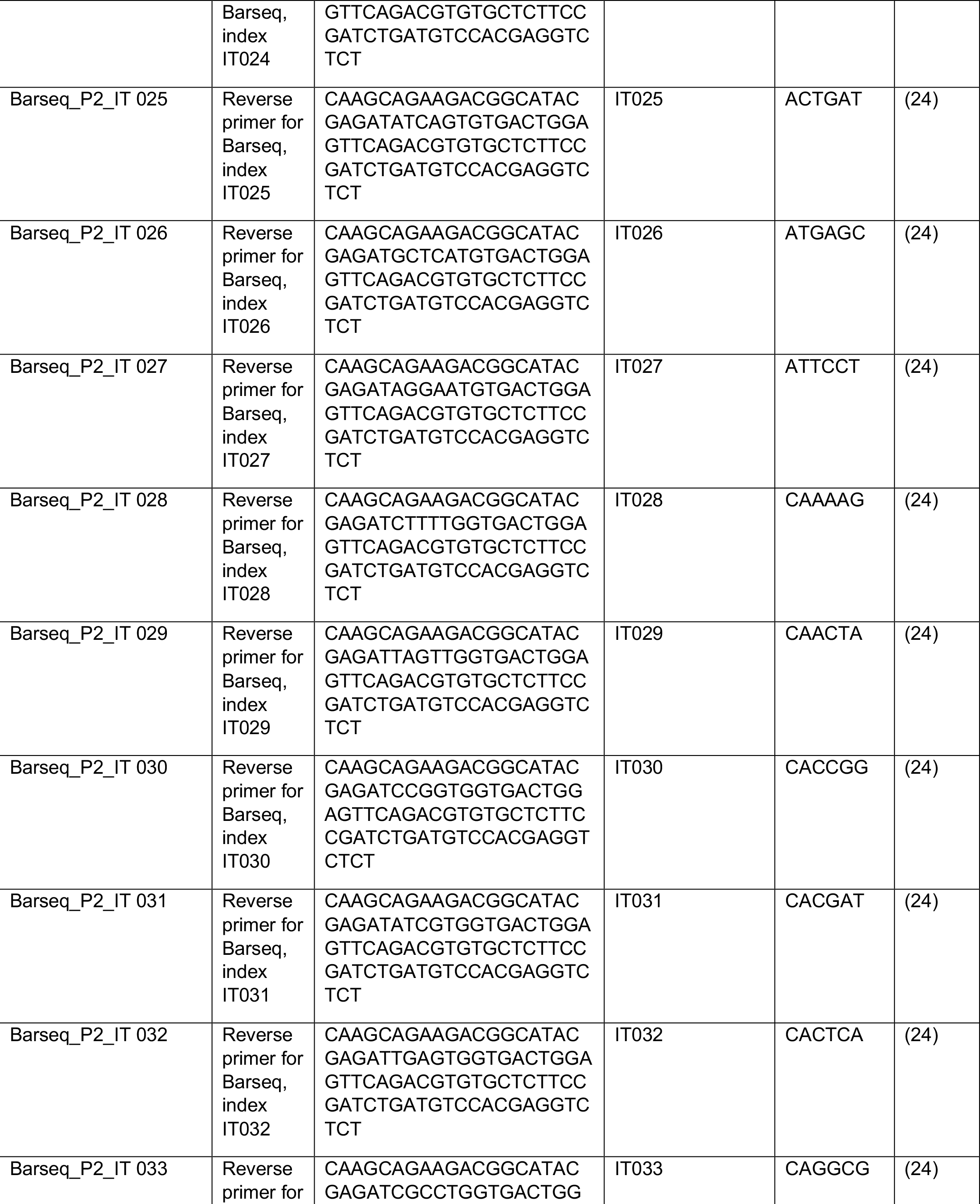
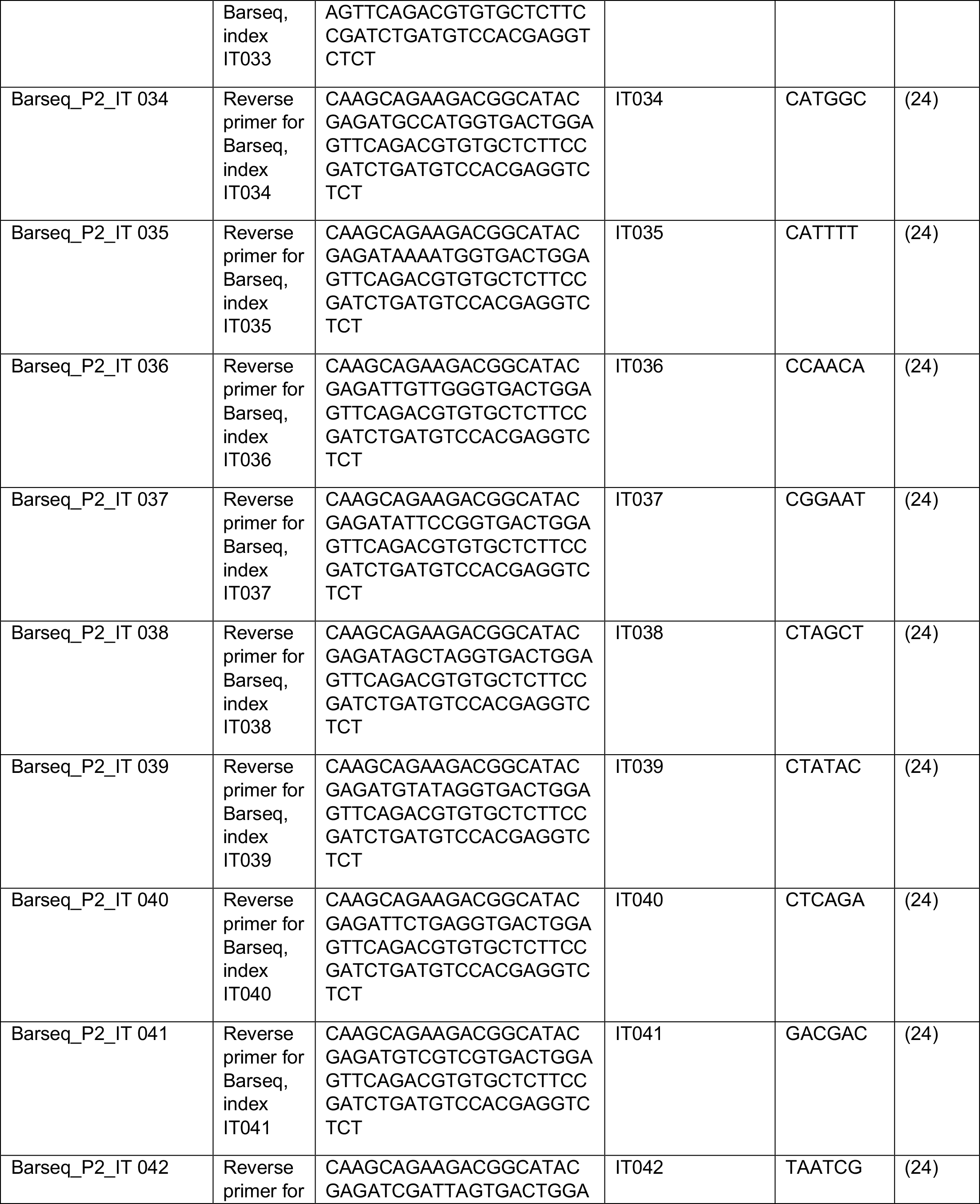
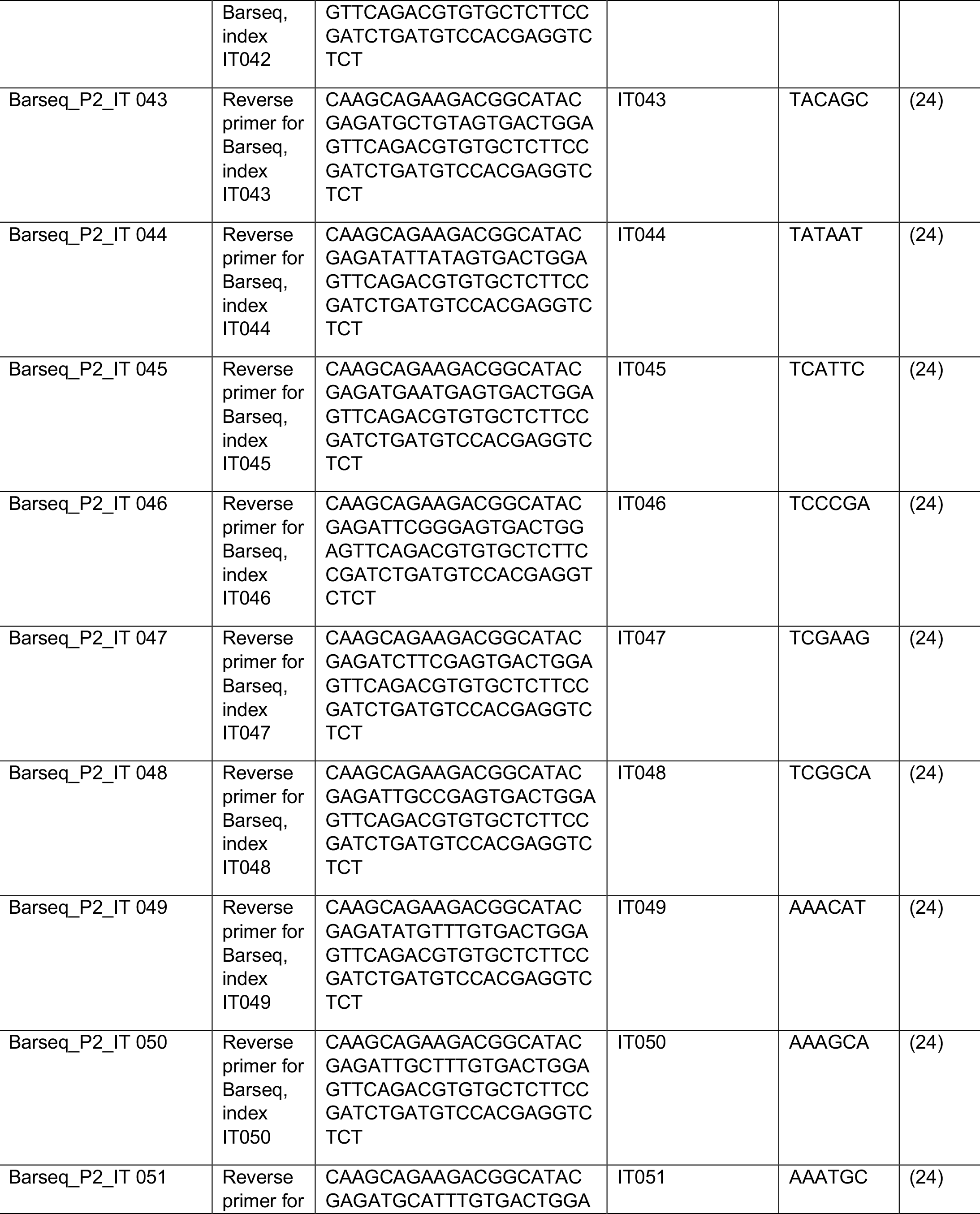
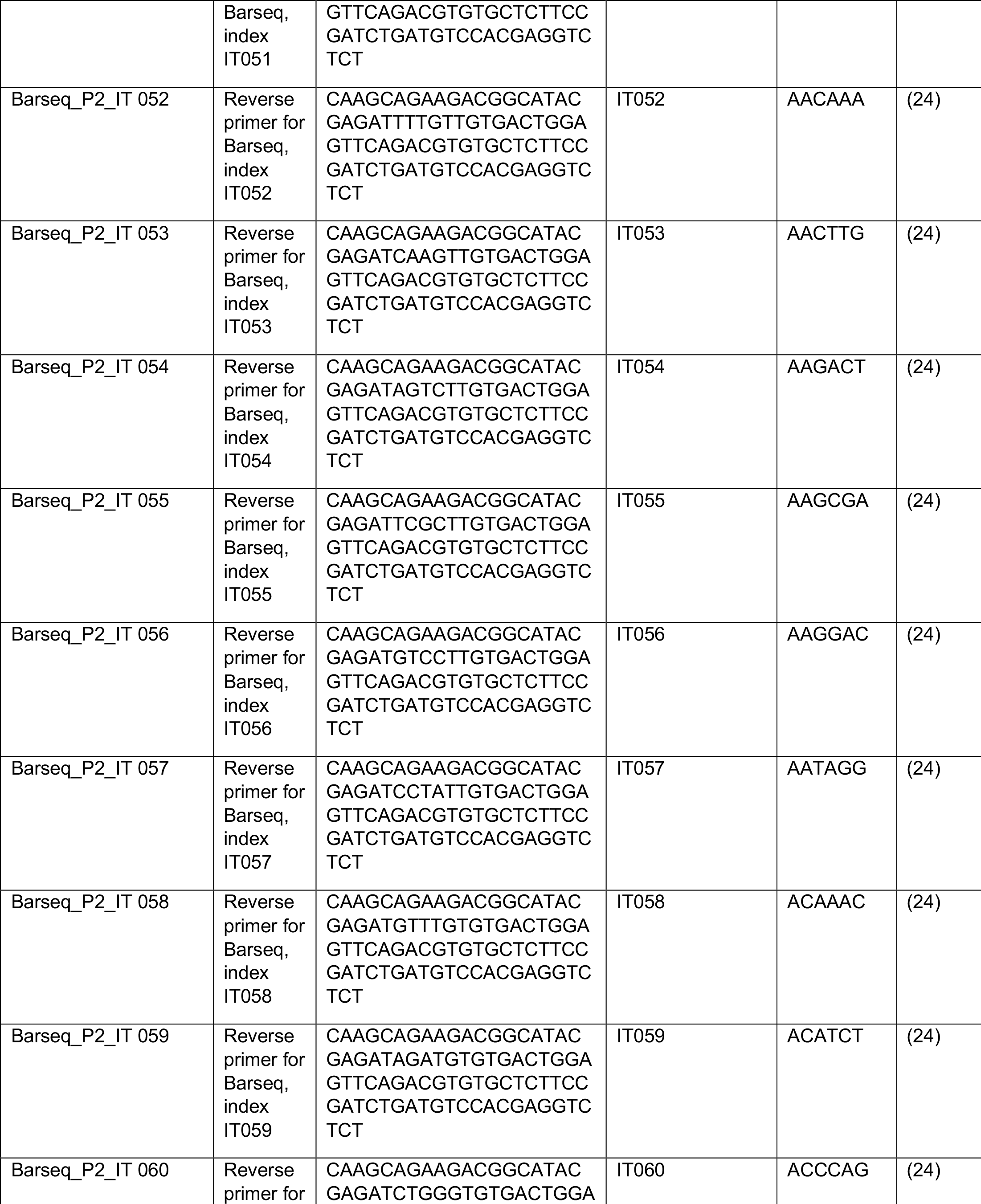
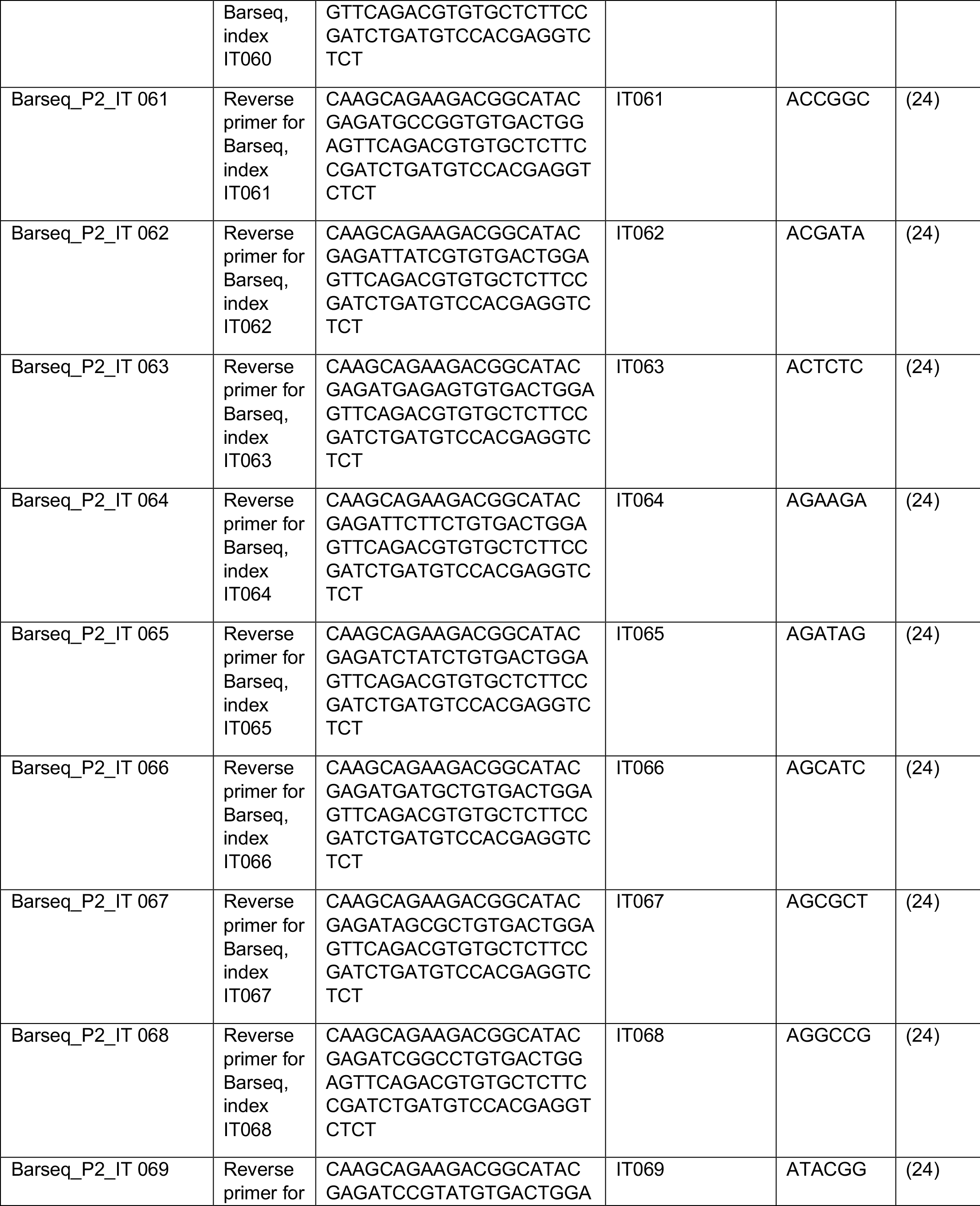
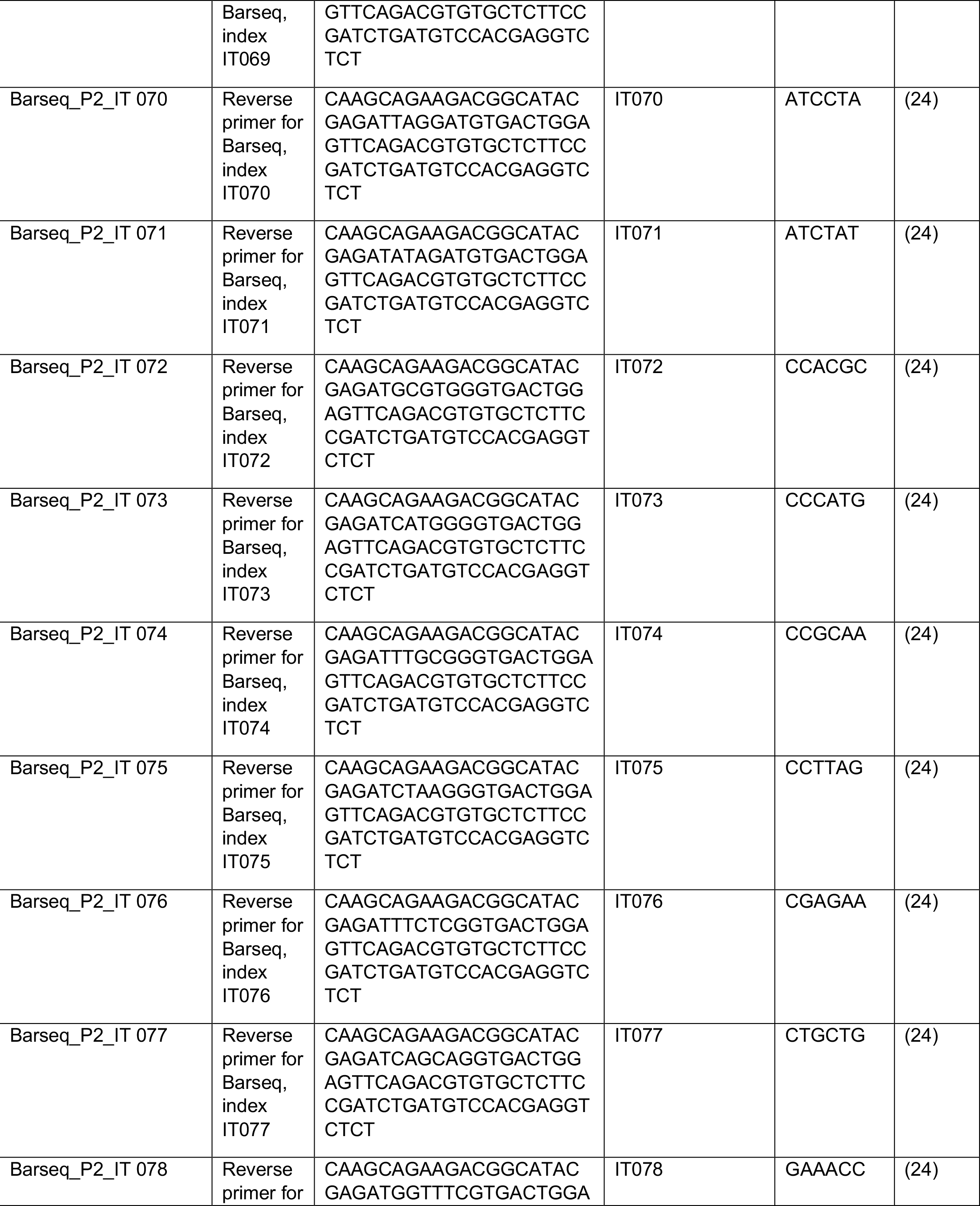
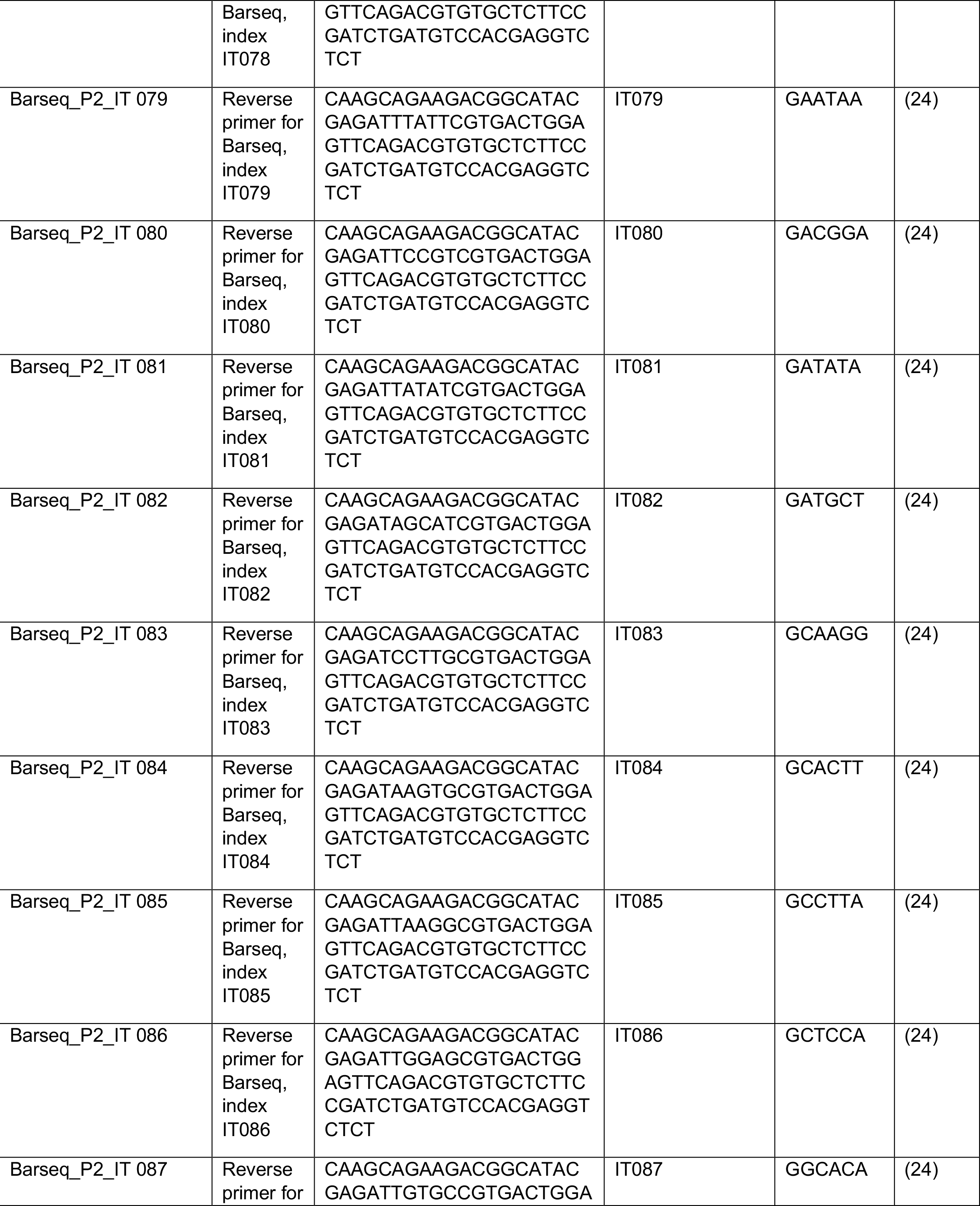
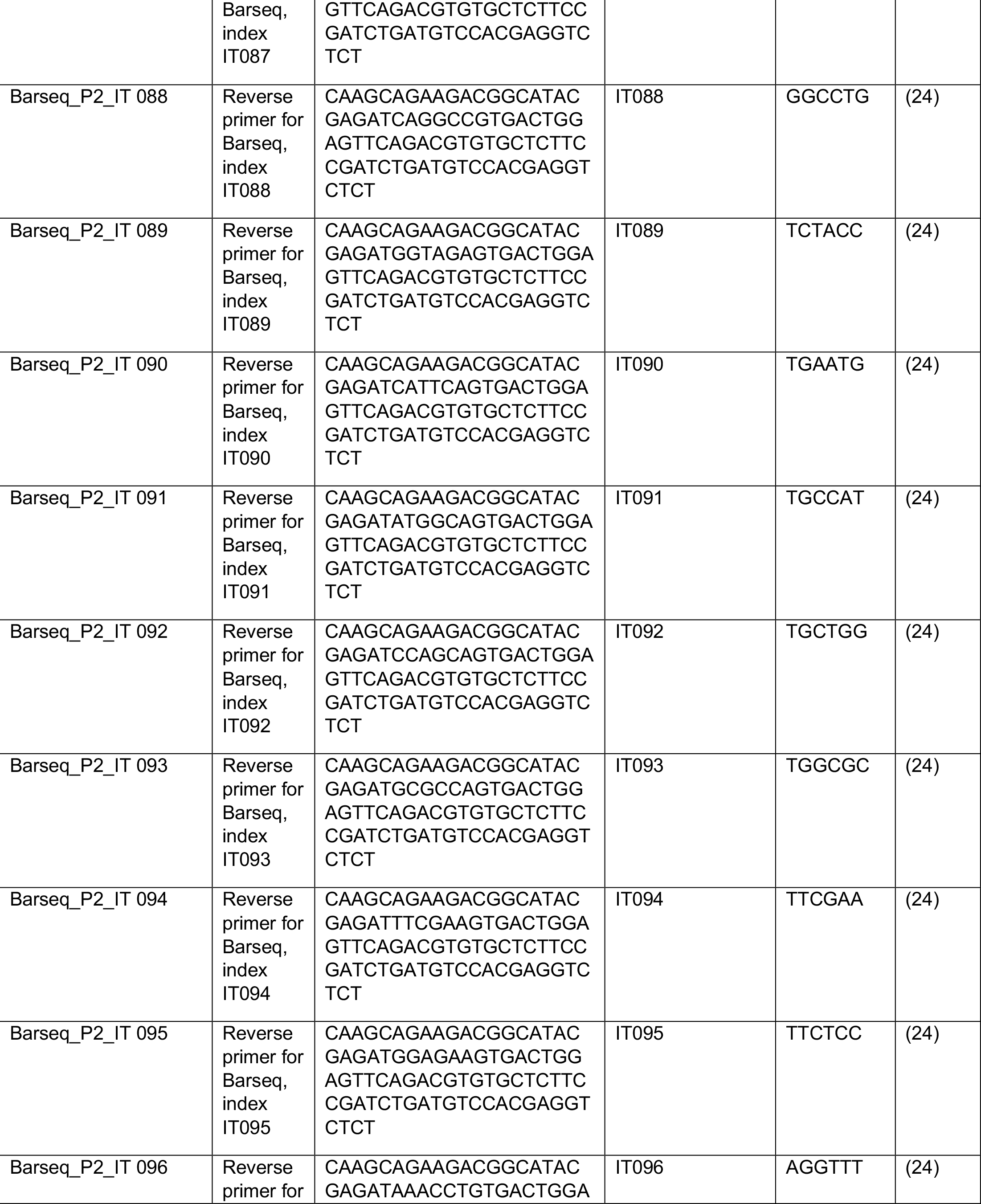
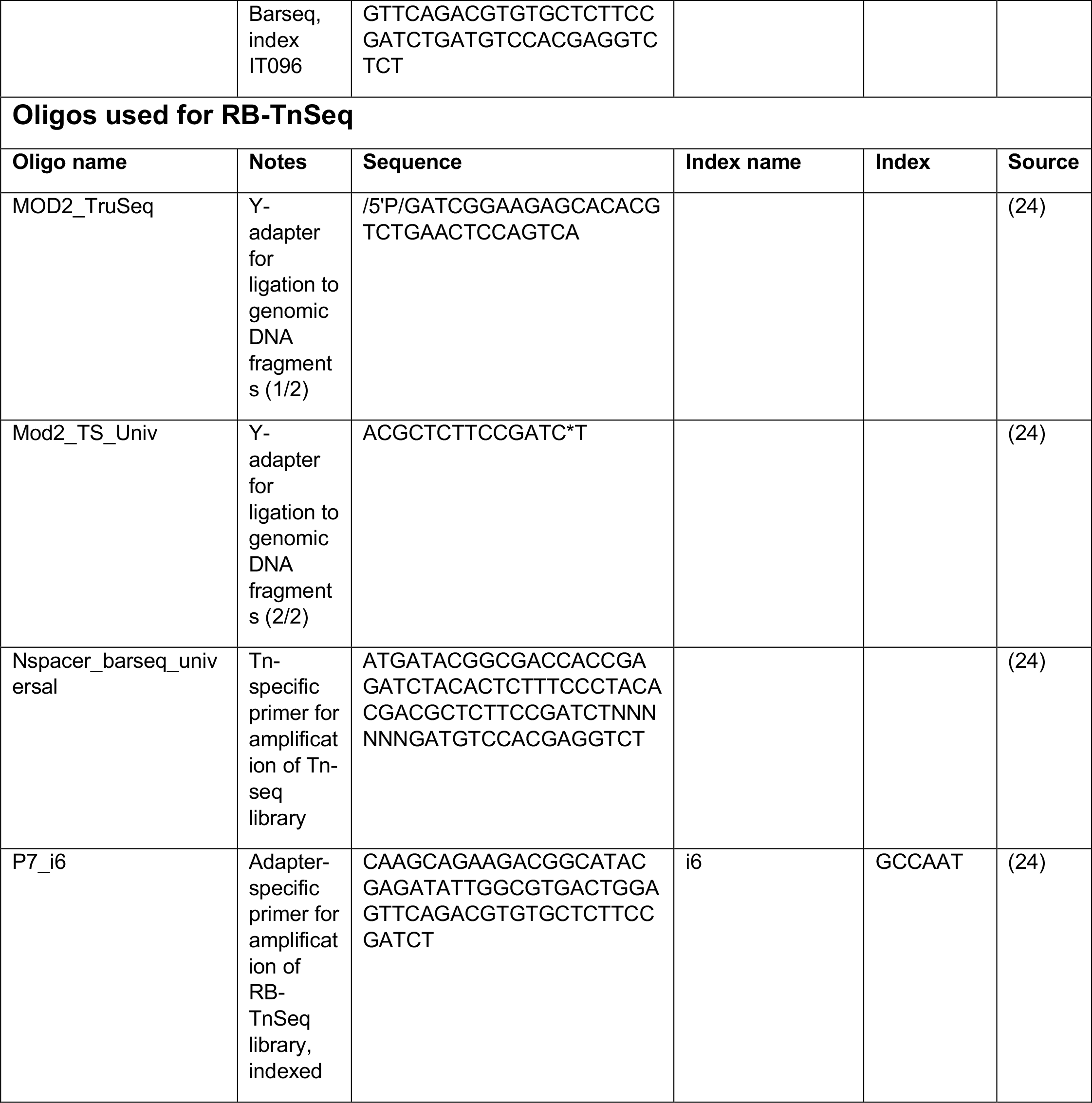
Strains and oligos used in this study.

**Table S2:** Program settings for re-arraying.

| Setting | Value |
| --- | --- |
| Volume | 40 µL |
| Pipetting tool | TS_300 |
| Transfer type | Pipette |
| Use filter tips | True |
| Change tips | Before each aspiration |
| Aspiration | Default |
| Dispensing | Dispense from defined height: 10.0 mm from bottom. No additional movement. |
| Mix before aspirating | False |
| Mix after dispensing | False |
| Standard liquid type | Protein |
| Change parameter | Yes (modified) |
| Aspiration speed | 7 mm/s |
| Dispensing speed | 7 mm/s |
| Blow delay | 0 ms |
| Blow speed | 11 mm/s |
| Immersion depth aspiration | -3 mm |
| Immersion depth dispense | 3 mm |
| Initial stroke | 0 % |
| Blow movement | 0 % |
| Prewetting | 0 cycles |

**Table S3:** Program settings for the inoculation of growth curves.

| Setting | Value |
| --- | --- |
| Volume | 2 µL |
| Pipetting tool: | TM_50_8 |
| Transfer type | Pipette |
| Use filter tips | Yes |
| Change tips | Before aspirating from a new source |
| Aspiration | Aspirate from defined height: 10.0 mm from bottom. No additional movement. |
| Dispensing | Dispense from defined height: 2.0 mm from bottom. No additional movement. |
| Mix before aspirating | No |
| Mix after dispensing | Yes |
| Standard liquid type | Protein |
| Change parameter | Yes (modified) |
| Aspiration speed | 11 mm/s |
| Dispensing speed | 11 mm/s |
| Blow delay | 0 ms |
| Blow speed | 11 mm/s |
| Immersion depth aspiration | -3 mm |
| Immersion depth dispense | 3 mm |
| Initial stroke | 0 % |
| Blow movement | 0 % |
| Prewetting | 0 cycles |
| Dispense mix: number of cycles | 1 |
| Dispense mix: speed | 11 mm/s |
| Dispense mix: volume | 40 µL |
| Dispense mix: fixed height | No |

